# Systematic Single Cell Pathway Analysis (SCPA) reveals novel pathways engaged during early T cell activation

**DOI:** 10.1101/2022.02.07.478807

**Authors:** Jack A. Bibby, Divyansh Agarwal, Tilo Freiwald, Natalia Kunz, Nicolas S. Merle, Erin E. West, Andre Larochelle, Fariba Chinian, Somabha Mukherjee, Behdad Afzali, Claudia Kemper, Nancy R. Zhang

## Abstract

Next generation sequencing technologies have revolutionized the study of T cell biology, capturing previously unrecognized diversity in cellular states and functions. Pathway analysis is a key analytical stage in the interpretation of such transcriptomic data, providing a powerful method for detecting alterations in important biological processes. Current pathway analysis tools are built on models developed for bulk-RNA sequencing, limiting their effectiveness when applied to more complex single cell RNA-sequencing (scRNA-seq) datasets. We recently developed a sensitive and distribution-free statistical framework for multisample distribution testing, which we implement here in the open-source R package Single Cell Pathway Analysis (SCPA). After demonstrating the effectiveness of SCPA over commonly used methods, we generate a scRNA-seq T cell dataset and characterize pathway activity over early cellular activation and between T cell populations. This revealed unexpected regulatory pathways in T cells, such as an intrinsic type I interferon system regulating T cell survival and a reliance on arachidonic acid metabolism throughout T cell activation. A systems level characterization of pathway activity in T cells across multiple human tissues also revealed alpha defensin expression as a hallmark of bone marrow derived T cells. Overall, our work here provides a widely applicable tool for single cell pathway analysis, and highlights unexpected regulatory mechanisms of T cells using a novel T cell dataset.

## Introduction

Dysregulation of T cell responses is a hallmark of a large spectrum human disorders, including autoimmunity, cancer and infectious diseases (Kumar et al., 2018). Dissecting the molecular events that underly T cell activation, lineage specification and effector function is therefore paramount to the therapeutic targeting of T cell-mediated disease. The advent of transcriptomics, and particularly single cell RNA sequencing (scRNA-seq), has provided the opportunity to link large transcriptional networks to specific cellular activation states and biological functions (Nayak and Hasija, 2021). Although gene level changes upon T cell activation and/or lineage induction have been well studied, systematic pathway level analyses of T cell populations are lacking. Given the power of pathway analysis to uncover specific biological processes in sequencing data, a systems level analysis of pathway activity applied to many T cell lineages and across stimulation conditions provides the opportunity to reveal novel and key regulatory pathways.

Many approaches have been developed that attempt to untangle the complexity of scRNA-seq data, such as methods for integration (Hie et al., 2019; Stuart et al., 2019; Tran et al., 2020; Welch et al., 2019), trajectory inference (Saelens et al., 2019; Street et al., 2018; Wolf et al., 2019), and dimensionality reduction (McInnes et al., 2008; van der Maaten and Hinton, 2008). This being said, biological pathway analysis methods are uniquely underdeveloped. Current approaches rely on models from bulk RNA-sequencing, wherein the central focus is on the quantification of signals from individual genes to generate an enrichment or overrepresentation score (Huang da et al., 2009; Kuleshov et al., 2016; Ma et al., 2020; Subramanian et al., 2005). These methods are based on the assumption that enrichment of a given gene set is the most meaningful statistic when understanding pathway importance. However, these approaches significantly under-utilize the information in the multivariate distribution that a pathway can exhibit in single cell data. Moreover, conventional pathway analysis methods rely on the input of a filtered list of differentially expressed genes (e.g. DAVID, Enrichr) and/or are all currently limited to two-sample comparisons (e.g. GSEA). This means that not only are large quantities of potentially relevant data discarded or ignored, but also tracking pathways over more complex experimental designs, such as multiple time points, is difficult to achieve in a robust way. Therefore, methods that can utilize the multivariate complexity and increasingly multisample design of scRNA-seq studies provide the potential for a more nuanced understanding of gene set behavior.

Here we present Single Cell Pathway Analysis (SCPA), an open-source R package for pathway analysis, and apply it to characterize pathway activity over early T cell activation. SCPA is built around a graph-based nonparametric statistical model (Mukherjee et al., 2020) that aims to fully capture the multivariate complexity of single cell data without imposing parametric assumptions on the gene expression distribution. This represents a fundamentally different approach to pathway analysis, whereby gene set activity or perturbation is primarily understood as a change in the multivariate distribution of a given pathway. To gain insights into gene set dynamics underlying early

T cell activation and differentiation, we generate a T cell scRNA-seq dataset across early activation and apply SCPA, uncovering unexpected pathway signatures across time and between T cell populations. This includes revealing an intrinsic IFNα pathway that maintains T cell survival upon stimulation, and demonstrating T cell reliance on arachidonic acid metabolism for effective cellular activation and cytokine production. We also perform a systems level characterization of pathway activity across multiple tissue sites to reveal tissue specific features of T cells. In carrying out this analysis, we highlight multiple features of SCPA, including classical two-sample comparisons, multisample pathway analysis, and tracking gene set perturbations over a pseudotime trajectory. Finally, we demonstrate the scalability of SCPA and its ability to comprehensively characterize pathway level changes across multiple conditions. Overall, we present a user-friendly and highly sensitive tool for pathway analysis in scRNA-seq data along with several new insights into the gene set changes driving early human T cell activation. All documentation and tutorials for SCPA can be found at https://jackbibby1.github.io/SCPA.

## Results

### Single Cell Pathway Analysis (SCPA) outperforms current pathway analysis tools

We recently developed a nonparametric graph-based statistical framework (Mukherjee *et al*., 2020) for comparing multivariate distributions in high dimensional data. Here we implement this framework in Single Cell Pathway Analysis (SCPA); an open-source R package for the analysis of pathway activity in scRNA-seq data. In brief, this statistic assesses the multivariate, joint distribution of a set of genes belonging to a given pathway to infer whether this pathway is differentially regulated across conditions (see methods). The stepwise outline for SCPA is shown in Figure 1A.

**Figure 1.**
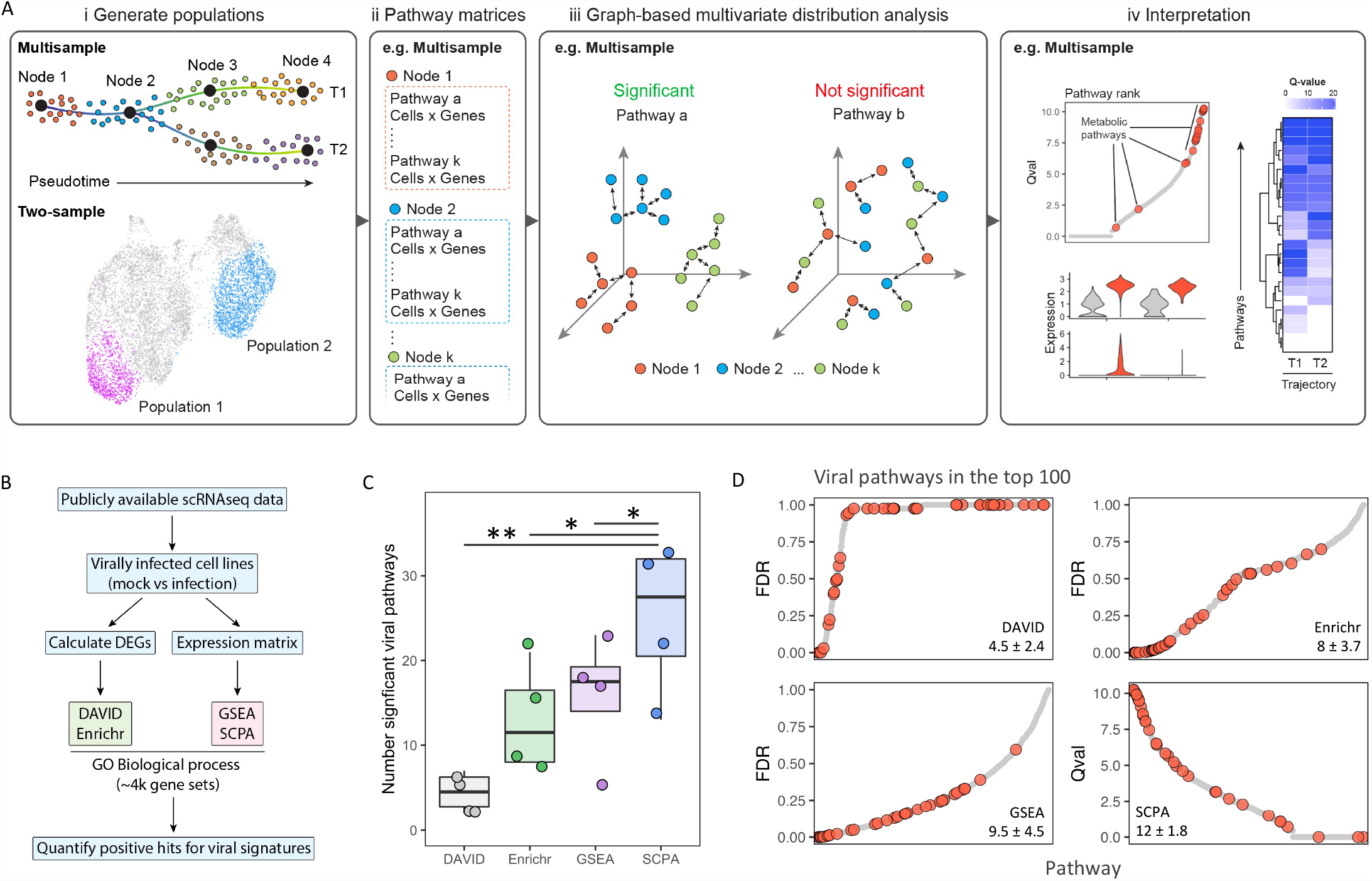
SCPA provides a sensitive and accurate reflection of pathway activity (A) Overview of the methodology implemented in the Single Cell Pathway Analysis (SCPA) R package. SCPA takes count matrices generated from the desired cell populations, generates nested matrices based on the genes of a pathway, and then performs graph based multivariate distribution analysis to assess pathway perturbation. The Q value output produced by SCPA can then be visualized for interpretation. (B) Overview of pathway analysis benchmarking. Briefly, publicly available scRNAseq data (GSE122031, GSE148729, GSE156760) that included cell lines infected with viruses were collated. Pathway analysis was then conducted comparing mock versus infected cell lines with either DAVID, Enrichr, GSEA, or SCPA, using ‘GO Biological Process’ gene sets. The number of significant viral pathways, and how many viral pathways are present in the top 100 pathways, were then compared across methods. (C) The number of viral pathways reaching significance when comparing mock to virally infected cells, across four publicly available datasets using the indicated method. (D) The number of viral pathways that rank in the top 100 pathways. Dot plot shows the rank of viral pathways relative to all GO Biological Process pathways across methods. The number below each method represents how many viral pathways were identified in the top 100 as mean ± SD. All viral pathways are shown in red, and non-viral pathways are shown in grey.

SCPA takes total normalized count matrices from two or more conditions (Figure 1Ai), preserving the complete dataset. We extract pathway matrices for any given number of pathways, based on manually curated or existing annotation databases (Figure 1Aii). Next, based on the joint distribution for all the genes of a given pathway, cells are paired based on optimal matching in the multidimensional space (Figure 1Aiii), where the number of dimensions is derived from the number of genes. Whether cell pairings occur between cells belonging to the same or different condition determines whether that pathway is differentially distributed or not. For instance, a high number of within-sample matches for a given condition suggests differential distribution of a pathway (left side Figure 1Aiii). Conversely, many inter-sample matches would suggest that a pathway is not differentially distributed across the groups or conditions (right side Figure 1Aiii). This analysis provides a statistic, here termed the Q value, which measures the size of distribution change for a given pathway and can be used for the ranking of pathways in order of biological relevance (Figure 1Aiv). This method therefore provides a fundamentally different definition of pathway activity when compared to current tools that typically rely on enrichment of overrepresentation of genes in a given pathway. By design, this method is robust to outliers and is shift- and scale-invariant, i.e. the Q value does not change if the expression values of all genes in a given pathway were scaled up or shifted by a constant factor.

Before using SCPA to analyze T cell biology, we first benchmarked the sensitivity and accuracy of SCPA against commonly used pathway analysis tools; notably DAVID and Enrichr that use differentially expressed genes as an input, and GSEA that uses total count matrices as an input (Figure 1B). We analyzed four publicly available scRNA-seq datasets generated from mock or virally infected – either influenza or SARS-CoV – cell lines. This allowed us to define positive controls for expected virus-induced gene signatures in a dataset, and then quantify the ability of different pathway analysis methods to detect these expected viral signatures. For the analysis, we used the GO Biological Process gene sets, given that they contain a large range of cellular pathways, and within this, a large range of viral signatures (see methods).

We first quantified how many viral related pathways were detected as differentially regulated using each method. We found that SCPA outperformed analytical tools that utilize lists of differentially expressed genes, detecting significantly more viral signatures in virally infected cells than DAVID and Enrichr. GSEA and SCPA were able to identify larger numbers of significant viral signatures, although SCPA significantly outperformed GSEA, identifying ∼30% more significant viral signatures (Figure 1C), demonstrating that SCPA is a highly sensitive method for detecting pathway perturbations in scRNA-seq data. In addition to quantifying the number of significantly altered viral pathways, we also asked where these viral pathways rank relative to other differentially regulated pathways in the analysis. Here, as a measure of methodological accuracy, we expected that viral pathways should rank highly, given that this was the only treatment variable across the two conditions (mock vs infected). To measure this, we assessed how many viral pathways were present within the top 100 significant pathways. Here, SCPA consistently identified a greater number of viral pathways present within the top 100 pathways relative to other methods (Figure 1D); on average, SCPA identified 12 viral pathways in the top 100, compared to 9.5, 8, and 4.5 for GSEA, EnrichR, and DAVID respectively.

In sum, we provide a novel package for pathway analysis that is specifically powered to detect pathway perturbations in single cell data, which is both highly sensitive and accurate. SCPA was able to detect more significant viral pathways, and these pathways were also more likely to be present in the top 100 significant pathways when compared to other methods.

### Single cell sequencing on sorted and stimulated T cell populations

We next utilized a systems level approach with SCPA to understand early signals required for the generation of effective human CD4^+^ and CD8^+^ T cell responses. We first purified human naïve and memory CD4^+^ and CD8^+^ T cells via magnetic bead enrichment and subsequent FACS sorting, and cells were then either left unstimulated, or stimulated for 12 or 24 hours via anti-CD3 and anti-CD28 antibodies (Supplementary Figure 1). scRNA-seq was then performed on each T cell population, capturing a total of over 40,000 live cells (Figure 2A).

**Figure 2.**
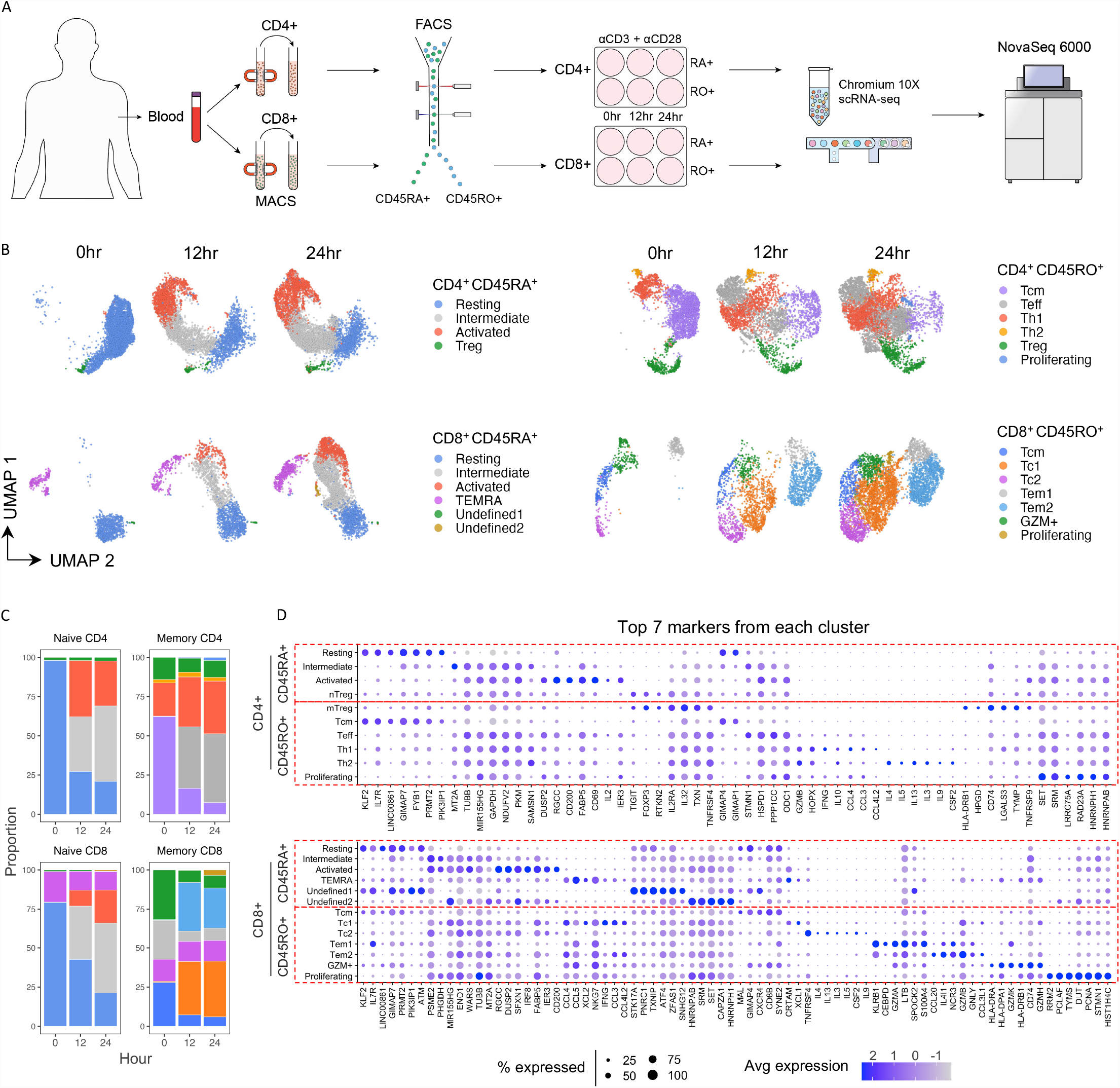
Single cell sequencing on sorted and stimulated T cell populations (A) Overview of experimental design for generation of the T cell scRNA-seq resource. CD4^+^ and CD8^+^ T cells were magnetically isolated from the peripheral blood in parallel, stained for CD45RA or CD45RO, FACS sorted to distinguish naïve from memory T cells, and left unstimulated, or stimulated with anti-CD3 and anti-CD28 for either 12 or 24 hours. Cells then underwent single cell sequencing and subsequent downstream analysis. (B) UMAP representations of T cell subtypes identified in the peripheral blood. Data were integrated separately across time points for each of the four cell types e.g. integration across naïve CD4^+^ T cells at 0, 12, 24 hours. Conditions were then split by time point for visualization. (C) Proportions of each identified cell type across each condition. Colors for bars are matched with colors in UMAP representations. (D) Dot plot representations of the markers from each cell cluster. Marker identification was done by combining data across stimulation time points for each cell type. An unbiased selection of markers was then generated by taking the top 7 genes sorted by false discovery rate from each population.

Within the scRNA-seq data generated here, we identified well defined T cell populations, both in the resting and activated states. All samples, populations, and time points identified in the sequencing are depicted in the UMAP projections (Figure 2B), proportional fractions of cell populations are shown in Figure 2C, and an unbiased selection of top 7 markers (as sorted by false discovery rate) from each cluster are shown in the corresponding dotplots (Figure 2D). All markers for each population can also be found in Supplementary Table 1.

Unsupervised clustering identified between 4 to 7 sub-populations within the different samples, which are indicated in the UMAP projections (Figures 2B-D). In the naïve populations, both CD4^+^ and CD8^+^ T cells contained a relatively homogenous resting population, expressing high levels of quiescence markers (*IL7R, KLF2*, and *PIK3IP1*). We also identified natural regulatory T cells (Tregs, expressing *FOXP3, IL2RA*, and *CTLA4*) and T effector memory cells re-expressing CD45RA (TEMRA, expressing *CCL3, CCL4, GZMB*, and *IFNG*) populations in the CD4^+^ and CD8^+^ samples, respectively. Naïve CD8^+^ T cells also included two small populations of unidentified cells. Upon stimulation, both CD4^+^ and CD8^+^ T cells generated two distinct populations corresponding to early and late phases of naïve T cell activation (left panels, Figure 2B). These data, in combination with trajectory inference modelling, showed that these activation phases included: 1) early expression of many guanylate-binding proteins alongside IRF1 and IRF4, and 2) subsequent expression of several heat shock proteins, a high number of metabolic genes, and effector molecules such as TNFα (Supplementary Figure 2).

As anticipated, circulating CD4^+^ and CD8^+^ T cell memory populations were less homogenous than circulating naive T cells, clustering into 6 and 7 sub-populations respectively. Clusters of effector cells were evident in resting and activated memory T cells. In memory CD4^+^ T cells, we identified T central memory (Tcm), T helper 1 (Th1), Th2, T cells with a high expression of CD69 and IL2 (here termed Teff), and Tregs, based on their canonical markers shown in Figure 2D. Tcm cells represented a large fraction (∼65%) of resting memory T cells, expressing markers very similar to resting naïve populations (*IL7R, KLF2*, and *TCF7*). Similar to CD45RA^+^ Tregs, CD45RO^+^ Tregs expressed high levels of *FOXP3*, but also effector molecules such as *IL32, HLA-DRB1, HLA*-*DRA*, and the HLA class II molecule *CD74*. Upon activation of memory CD4^+^ T cells, we observed a large proportional expansion of Teff cells, expressing high levels of *IL2* and *CD69*. In the differentiated effector populations, Th1 cells showed high expression of *IFNG, IL10, CCL3, CCL4*, and *GZMB* upon activation, whereas Th2 cells expressed high levels of *IL3, IL4, IL5, IL9, IL10*, and *GATA3*. In the memory CD8^+^ T cell compartment, we identified Tcm cells expressing quiescence markers such as *KLF2*. Similar to CD4^+^ T cells, we identified effector populations including Tem1/2 cells expressing *GZM* and *LT* genes, Type 1 CD8^+^ T cells (Tc1) expressing *XCL1, XCL2, IFNG, CCL3*, and *CCL4*, and Tc2 cells expressing *IL3, IL4, IL5, IL9*, and *IL13*. We also identified a population (here termed GZM^+^) that expressed high levels of all *GZM* genes (Supplementary Figure 3), and also multiple HLA molecules. Finally, and similar to memory CD4^+^ T cells, we detected a cell population dominated by proliferation markers at 24 hours post stimulation.

Overall, this analysis identifies well resolved T cell populations, with cell types being stratified by canonical markers. These data therefore provided a resource for characterization of pathway level transcriptomic signatures through early activation, and across a range of human CD4^+^ and CD8^+^ T cell populations, as detailed below.

### Intrinsic IFNα signaling as a regulator of T cell survival

We next applied SCPA to map gene set dynamics throughout T cell activation. To gain a global view of pathway activity within each of the four major T cell populations, we first used the multisample capability of SCPA – a technique not possible using current methods – to characterize pathway distributions over the three time points simultaneously for each sorted population (0, 12, and 24 hours for naïve CD4^+^, memory CD4^+^, naïve CD8^+^, and memory CD8^+^ populations, Figure 3A). We used a comprehensive gene set list that included all pathways from the Hallmark, Reactome, and KEGG databases from MSigDB, comprising 1790 pathways in total (Supplementary Table 2). Utilizing this gene set list, we identified a set of ‘core pathways’ defined as common gene sets showing significant activity in response to stimulation across all cell types (Figure 3B). After categorizing these gene sets into broad classes, the most abundant pathway classes were, as expected, those involved in response to stimuli, including cell signaling, cell cycle, cytokine response, metabolism, and transcription (Figure 3B). In addition to this core signature, we also observed a cluster of pathways that typically showed greater activity in memory T cells versus naïve populations upon stimulation, such as unfolded protein response and autophagy (Supplementary Figure 4). Further, certain pathways showed little or no change in distribution upon activation, including pathways not expected to change, such as digestion (Supplementary Figure 4). Interestingly, across pathways, memory cells showed a larger magnitude of change in their pathway distribution upon activation, suggesting that memory populations undergo a more extensive remodeling of their transcriptional profile, per pathway, relative to naïve cells (Figure 3C). Overall, we validate the multisample capability of SCPA in accurately recapitulating known core pathway signatures of T cell activation over time.

**Figure 3.**
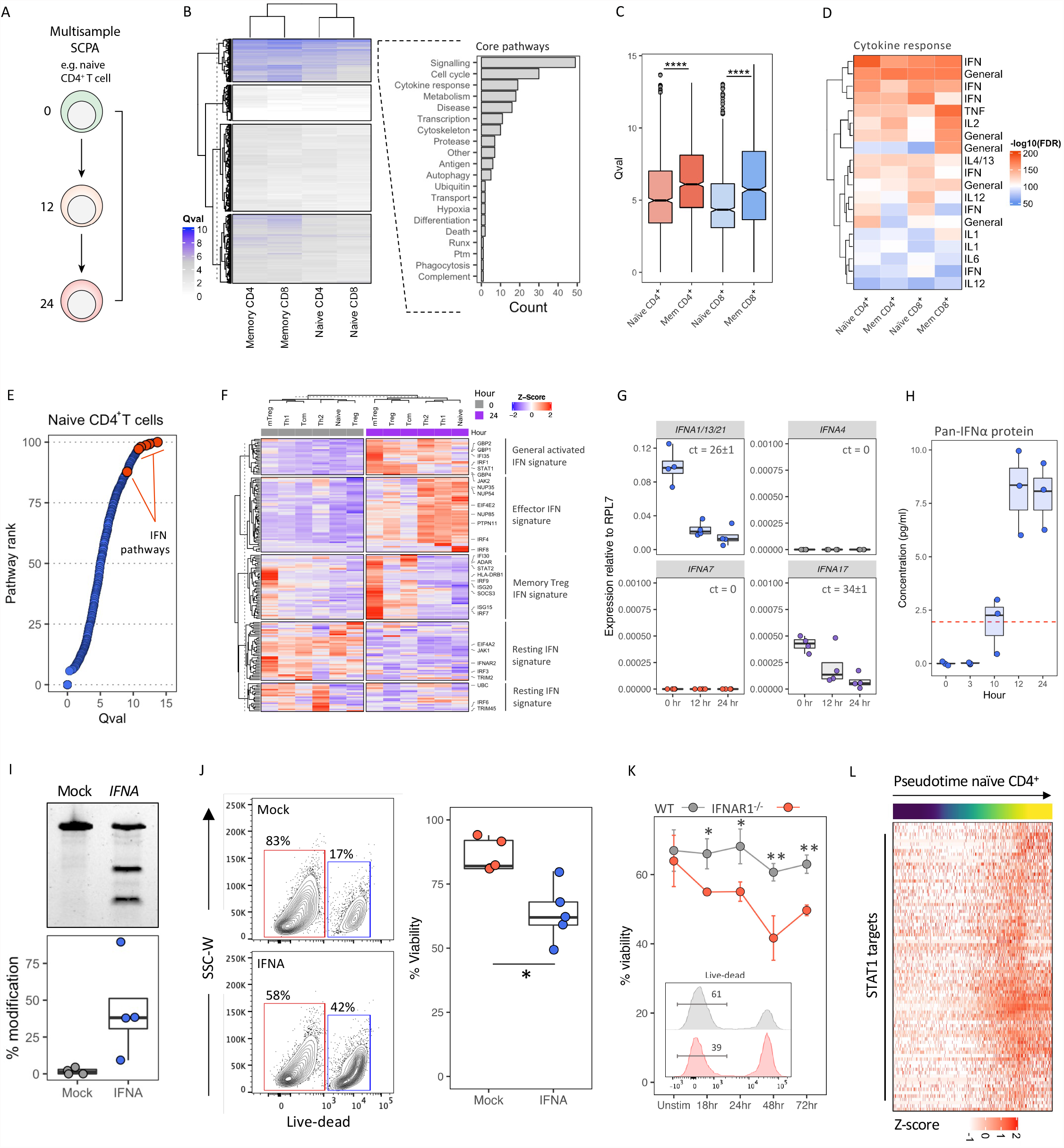
Intrinsic IFNα signaling as a key regulator of T cell survival (A) Schematic representation of the SCPA analysis conducted across time points for each sorted T cell population. Cells were split across time and SCPA was used to conduct a multisample analysis across the three time points simultaneously. (B) Heatmap representation of pathway perturbations generated from the comparisons outlined in (A), with classifications of core pathways into broad categories. The pathways from the topmost cluster (k-means clustering) were manually categorized into broad pathway classes, and the frequency of each class is visualized in the bar plot. (C) Boxplots representing the extent of distribution changes, per pathway, for all 1790 pathways across cell types and stimulation. (D) Heatmap showing FDR values of specific cytokines within the ‘cytokine response’ class derived from core pathways. ‘General’ refers to cytokine response gene sets that do not mention a specific cytokine e.g. ‘cytokine signaling’. (E) Ranking of all pathway Qvals in naïve CD4^+^ T cells across activation. Interferon (IFN) pathways are highlighted in red (F) Heatmap representation of interferon response gene expression with genes taken from the ‘Reactome interferon signaling’ gene set. (G) qPCR for IFNA genes in CD4^+^ T cells after stimulation with αCD3+ αCD28 for the indicated time with 0hr representing unstimulated cells. Data are relative to RPL7 as an internal reference, calculated as 2^-Δct^. Ct values represent the mean over 4 donors ± sd in the 0hr condition (H) IFNα measured in the supernatant of CD4^+^ T cells after stimulation with αCD3+ αCD28 for the indicated time. 0hr represents unstimulated cells, and red dotted line represents the detection limit (I-J) CRISPR-cas9 mediated deletion of IFNα in CD4^+^ T cells, showing IFN editing efficiency in (I) and flow cytometry analysis of cell viability after IFNα knockdown in CD4^+^ T cells (J). CD4^+^ T cells were isolated, and CRISPR mediated knockdown of IFNα was performed. Cells were then stimulated with αCD3+ αCD28 for 24 hours before analysis by flow cytometry. (K) Viability staining of splenic CD4^+^ T cells from wild type (WT) or Ifnar1^-/-^ mice. CD4^+^ T cells were purified and stimulated with anti-CD3 + anti-CD28 for the indicated time before assessment of viability. Embedded panel shows representative live-dead staining taken from the 48hr time point, n = 3 mice per genotype (L) Heatmap representation of STAT1 target gene expression over naïve CD4^+^ T cell activation. STAT1 targets, taken from the transcription factor targets available on MSigDB were plotted over a trajectory of naïve CD4+ T cell activation. Trajectory inference model calculated using slingshot is shown in supplementary figure 2A.

To more deeply understand these global profiles, we next looked for enrichment of specific signatures in the core pathways. As cell signaling and cell cycle events have been well characterized upon T cell activation, we chose to investigate cytokine response signatures. Within this pathway class we noted a presence of interferon (IFN) response gene sets across all four naïve and memory CD4^+^ and CD8^+^ T cell populations (Figure 3D). To further explore this, we focused on CD4^+^ T cells to assess if a particular cell type was driving this interferon response signature, and whether this pathway was being positively or negatively regulated upon activation. We used SCPA to conduct a two-sample comparison of each cell type, comparing 0hr to 24hr activated populations identified in the UMAP clustering (Figure 2B). Unexpectedly, IFN response genes were broadly enriched after activation in all CD4^+^ T cell populations (Supplementary Figure 5A). Taking naïve CD4^+^ T cells as an example, IFN response pathways also ranked amongst the most significantly altered pathways upon T cell activation (Figure 3E), suggesting that IFN signaling is a central component of T cell activation.

However, we observed that T cells induce a distinct IFN response signature, which is dependent on cell lineage. Most CD4^+^ T cells show a broadly similar profile in their resting state with only memory Tregs expressing a slightly altered signature. Cellular activation induced a divergence of this signature, with significant differences between memory Treg and non-Treg populations. For example, Tregs show enrichment of IFN response genes such as *ISG15, ISG20, IFI30*, and *SOCS3*, whereas non-Treg populations in the naïve and memory pool show enrichment of *IRF4, PTPN11*, and several *NUP* genes (Figure 3F). These data suggest that, even though all CD4^+^ T cells induce an interferon response signature upon activation, these signatures diverge depending on Treg versus non-Treg cell identity.

As these were sorted T cell populations, maintained in media without polarizing cytokines, and naïve CD4^+^ T cells do not produce type II *IFNG* transcript or IFNγ protein (Supplementary Figure 5B-C), these data suggested possible involvement of a cell-intrinsic type I interferon signaling. The significant overlap of genes between type I and type II response gene sets makes it difficult to define the exact interferon subtype driving the observed signature, but the above analysis with SCPA did reveal type I interferon signatures in both naïve and memory T cells (Supplementary Figure 5D). Although it is known that type I IFNs, IFNα and IFNβ, regulate CD4^+^ and CD8^+^ T cell cytokine production, proliferation, survival, and/or migration (Crouse et al., 2015; Marrack et al., 1999), T cells themselves are not considered to be a source of type I IFNs. Rather, it is thought that type I IFNs are largely derived from innate immune lineages during viral infections (Ivashkiv and Donlin, 2014). Therefore, we explored the possibility that T cells may produce their own type I IFN upon stimulation. Likely given the relatively low sensitivity of scRNA-seq, we did not detect *IFNA* expression in our dataset.

We therefore analyzed three RNA-seq/microarray datasets from the literature for expression of *IFNA* genes in CD4^+^ T cells either *ex vivo* or after *in vitro* activation. Here we observed low but consistent levels of *IFNA* expression in both resting and activated T cells, with the most consistent expression of *IFNA4, IFNA7, IFNA17*, and *IFNA21* across datasets (Supplementary Figure 5E). Given the increased sensitivity of qPCR over RNA-sequencing methods, we next measured expression of these four *IFNA* genes upon T cell activation using qPCR. We observed robust expression of *IFNA1/13/21* in CD4^+^ T cells and low levels of *IFNA17*, whereas we detected little or no expression of *IFNA4, IFNA7* (Figure 3G). Additionally, and in agreement with our qPCR data, we noted that CD4^+^ T cells secrete low levels of IFNα protein upon stimulation at around 12-24 hours post activation with anti-CD3 and anti-CD28 (Figure 3H, Supplementary Figure 6E).

Given that we observed IFNα production by CD4+ T cells, we next asked if an absence of T cell-derived IFNα resulted in an altered T cell phenotype. We therefore used CRISPR-Cas9 mediated deletion of IFNα in purified CD4^+^ T cells, targeting multiple *IFNA* genes, afforded by the high sequence similarity (see methods, Supplementary Figure 5G-H), demonstrating good editing efficiency (Figure 3I). Deletion of IFNα significantly reduced T cell viability upon stimulation, when compared to a mock control (Figure 3J). In agreement with this observation, purified CD4^+^ T cells from Ifnar^-/-^ mice – in lieu of Ifna^-/-^ mice not existing – showed a similar reduction in cell viability after activation *in vitro* with anti-CD3 + anti-CD28 (Figure 3K). Finally, IFNAR1 signaling engages downstream STAT1 signaling, and in a pseudotime trajectory analysis of naive CD4^+^ T cell activation, we observed a large increase in the expression of STAT1 responsive genes (Figure 3L, Supplementary Figure 5I), corresponding to ‘intermediate’ and ‘activated’ cells states identified in the UMAP representations (Figure 2B), and also correlated with the upregulation of *IFNAR1/2* by T cells (Supplementary Figure 5J). Together these data indicate that T cells require IFNα to maintain survival upon activation and that they possess the ability to produce IFNα in an autocrine/paracrine fashion.

### Mapping metabolic reprogramming upon T cell activation

After cytokine response pathways, our analysis with SCPA demonstrated that the gene set class with the next largest distribution change was that of metabolism (Figure 3B). Although context-dependent metabolic programs have been linked to specific T cell functions, a systematic assessment of transcriptional metabolic reprogramming in T cells has yet to be done. We therefore sought to characterize the landscape of transcriptional metabolic reprogramming upon T cell activation across comprehensive list of metabolic pathways.

SCPA’s unique capability to analyze multiple groups/conditions simultaneously lends itself to novel analyses of pseudotime trajectories with multiple intermediate populations. To exhibit this, we first reconstructed the trajectory of naïve CD4^+^ T cell activation, using slingshot (Street *et al*., 2018), to recapitulate the progression of these cells through activation phases. To generate distinct populations across activation, we split this trajectory across pseudotime into multiple nodes (using milestones as explained in methods, Figure 4A). Using a manually curated list of 243 metabolic pathways (gathered from Hallmark, KEGG and Reactome databases, Supplementary Table 3), we conducted a multisample pathway analysis across the pseudotime trajectory to identify metabolic pathways showing significant changes in distribution through naïve CD4^+^ T cell activation. Overall, we saw a gradation in the transcriptional changes of metabolic pathways, from pathways with little change to those with large changes in distribution (Figure 4B). The largest alterations were evident glycolytic, oxidative phosphorylation (OXPHOS), and amino acid metabolism pathways (Figure 4B, n.b multiple glycolysis and OXPHOS gene sets from different databases were present in the top 10 pathways, so only the topmost significant are annotated), indicating that these pathways are critically regulated at the transcriptional level in T cells. Indeed, as expected, glycolysis and OXPHOS were both enhanced following naïve and memory CD4^+^ T activation *in vitro* (Supplementary Figure 6A-B), consistent with current literature (Chapman et al., 2020). Interestingly, metabolic change, both relating to magnitude and pathway specificity in naïve T cells, was very similar to that seen upon memory T cell activation, suggesting a similar utilization of metabolic pathways in both populations (Supplementary Figure 6C). Conversely, perturbations in activity of other pathways, such as linoleic acid and nitrogen metabolism, were comparatively modest following T cell activation, possibly indicating that these pathways play a less significant role in the biology of activated T cells or that they are not regulated principally through transcription. Indeed, the pathway expression of linoleic acid metabolism over pseudotime is substantially different from glycolysis (Figure 4C). Additionally, we discovered several metabolic pathways that have so far not been linked as intrinsic regulators of T cell biology, such as propanoate metabolism and arachidonic acid metabolism (Figure 4B). Although undergoing smaller changes relative to pathways such as glycolysis and OXPHOS, these pathways showed statistically significant changes in pathway perturbation, possibly suggestive an important role in T cell function.

**Figure 4.**
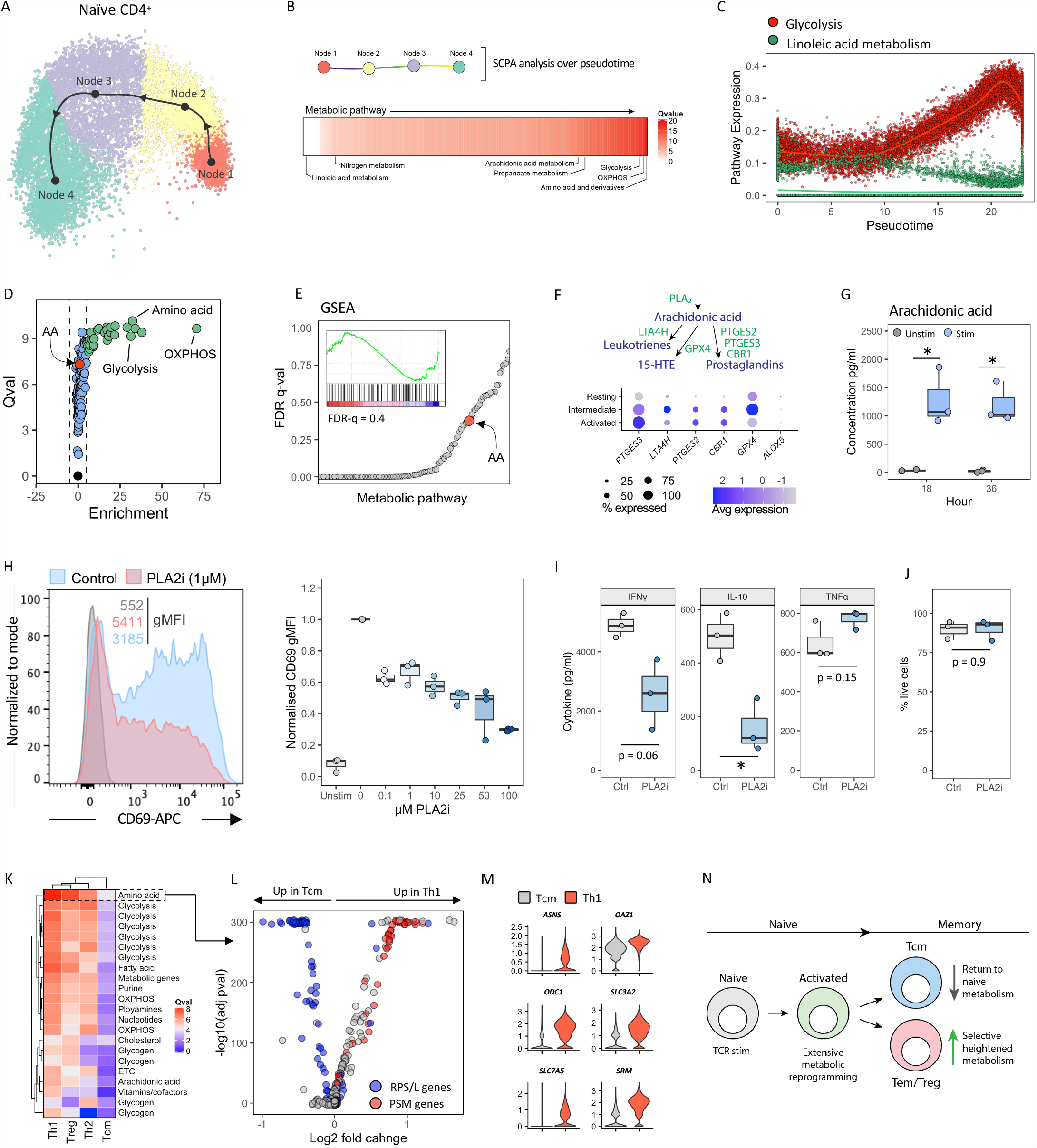
Arachidonic acid metabolism regulates CD4+ T cell activation (A) Trajectory analysis of naïve CD4^+^ T cell activation. Naïve CD4^+^ T cells were subjected to trajectory inference modelling using slingshot, and subsequently split into nodes across the trajectory using dyno (see methods). (B) SCPA analysis outline (above) and output (below) using a manually curated list of metabolic pathways across pseudotime populations. 186 manually curated metabolic pathways were used with SCPA to categorize metabolic reprogramming across the four identified activation phases in native CD4^+^ T cell activation. (CA) Mean pathway expression of glycolysis and linoleic acid metabolism over naïve CD4^+^ T cell activation. Mean gene expression for ‘Hallmark glycolysis’ and ‘KEGG linoleic acid metabolism’ were plotted against pseudotime values calculated in (A). (D) Volcano plot showing Qval output from SCPA plotted against pathway enrichment, measured as mean pathway change when comparing 0 to 24 hours conditions. Black points show non significant pathways, blue points show significant pathways with no enrichment, and green points show significant pathways that also show enrichment. Arachidonic acid metabolism (AA) is highlighted in red (E) Gene set enrichment analysis (GSEA) across T cell activation, using metabolic pathways when comparing 0 to 24 hours conditions. Dot plot shows ranking of metabolic pathways by FDR q-val, with arachidonic acid metabolism highlighted in red. Embedded plot shows enrichment plot for arachidonic acid metabolism. (F) Outline of key enzymes of the arachidonic acid metabolism pathway and their expression across activation. (G) Arachidonic acid production measured in the supernatants of purified CD4^+^ T cells after stimulation with anti-CD3+28 antibodies (H) CD69 expression in CD4^+^ T cells after inhibition of a PLA2 using pyrrophenone at varying concentrations. Representative plot taken from 1μM. (I-J) (I-J) Cytokine expression (I) and cell viability (J) after anti-CD3+28 stimulation with PLA2 inhibition (1μM) in CD4+ T cells (K) Heatmap of Qvals generated by SCPA after comparing the indicated resting CD4^+^ memory T cell population with resting naïve CD4^+^ T cells (L-M) Volcano plot of amino acid metabolism genes compared between Tcm and Th1 cells. Ribosomal (RPS/L) and proteasomal (PSM) genes are highlighted in blue and red respectively. (N) Schematic summary of T cell metabolism over cellular differentiation

As an internal control of the pseudotime pathway analysis, we correlated the results above with a more classical 2-sample comparison using our real time data. We compared 0hr resting cells, to either 12hr intermediate or 24hr activated naïve CD4^+^ T cell populations (populations derived from the UMAP in Figure 2B). In line with our pseudotime trajectory pathway analysis (Figure 4B), OXPHOS, glycolysis, and amino acid metabolism pathways were highly significant and also positively enriched in activated cells (Figure 4D, Supplementary Figure 6D). Interestingly, and an important unique aspect of our tool, many pathways showed significant changes in multivariate distribution, but no overall enrichment (Figure 4D, blue dots). Thus, these pathways show changes in transcriptional regulation detected by SCPA, that are independent of overall pathway enrichment. One such highly significant pathway showing this trend, outside of metabolic pathways already defined as important to T cell function, was arachidonic acid metabolism. We compared this finding to GSEA to see if such a signature would be missed using classical approaches, and indeed, given the lack of pathway enrichment, arachidonic acid metabolism was not significant, ranking poorly at both 12 (FDR q-val = 0.7) and 24 (FDR q-val = 0.4) hours after stimulation (Figure 4E, Supplementary Figure 6E). Given that the intrinsic regulation of arachidonic acid metabolism has not been described in T cells, we sought to confirm our finding with exploratory functional experiments. We noted upregulation of key enzymes involved in the generation of arachidonic acid and downstream metabolites upon T cell activation (Figure 4F), and found that T cells secrete arachidonic acid upon activation (Figure 4G). To understand the functional consequences of arachidonic acid synthesis in T cells, we used a Phospholipase A2 (PLA2) inhibitor, pyrrophenone, which blocks arachidonic acid synthesis from phospholipids. PLA2 inhibition resulted in a blunted upregulation of CD69 (Figure 4H), suggestive of decreased T cell activation. Furthermore, PLA2 antagonism resulted in decreased expression of IFNγ and IL-10, but showed no effect on TNFα production or cell viability (Figure 4I-J). Overall, these data demonstrate the benefit of understanding pathway activity in terms of changes in multivariate distribution in contrast to enrichment, and we validate this approach by showing the importance of PLA2 and downstream arachidonic acid-derived metabolites for T cell activation, effector function, and production of selected cytokines.

Having analyzed the transcriptional regulation of metabolic pathways upon activation, we next decided to investigate how metabolic pathways are maintained across CD4^+^ T cell differentiation. Although it is known that naïve and memory CD4^+^ T cells maintain different standards of glycolysis and OXPHOS (Dimeloe et al., 2016; Gubser et al., 2013), a more systematic approach into the landscape of metabolic gene expression, or how memory T cell populations differ, is unknown. For example, whether transcriptional regulation of metabolic pathways among Th1, Th2, and Tcm cells differs remains uncharacterized. We therefore performed a systematic pathway analysis across resting memory T cell populations (Th1, Th2, Treg, and Tcm) focusing on metabolic pathways, and considering naïve resting CD4^+^ T cells as a comparative baseline (Figure 4K). Here we observed two prominent features. First, we noted an extensive perturbation of metabolic pathways outside of glycolysis and OXPHOS in resting memory T cells, including pathways regulating amino acid metabolism, fatty acid metabolism, nucleotide synthesis, purine metabolism, cholesterol metabolism, glycogen synthesis, and arachidonic acid metabolism. This amounted to 0-15% of all annotated metabolic pathways being differentially regulated in resting memory populations versus resting naïve T cells (Supplementary Figure 6F). Second, we observed substantial differences between resting memory T cell subtypes, specifically effector subsets (Th1, Th2, and Tregs) and Tcm cells, with Tcm cells showing a broadly similar profile to resting naïve T cells (Figure 4K). Highlighting specifics, looking at topmost differentially regulated pathway, amino acid metabolism, we saw that a defining feature of Tcm vs Th1 cells is their relative expression of ribosomal versus proteasomal genes (Figure 4L). Here Tcm cells were defined by relatively high expression of ribosomal subunit (RPL/RPS) genes whereas Th1 cells showed higher levels of proteosome subunit (PSM) genes. Furthermore Th1 cells were characterized by higher expression of genes regulating polyamine synthesis (*OAZ1, ODC1, SRM*), asparagine synthesis (ASNS), and amino acid transport (*SLC3A2, SLC7A5*), suggesting that Tcm and Th1 cells differ significantly in their utilization and synthesis of amino acids (Figure 4M). Taken together, these data demonstrate that quiescent Th1, Th2, and Treg populations differ significantly in their transcriptional regulation across a wide range of metabolic pathways when compared to Tcm and resting naïve CD4^+^ T cells (Figure 4N).

Overall, these data demonstrate the multisample pseudotime capability of SCPA and highlight the power of a systematic approach identify new pathways underlying metabolic regulation of cellular activity.

### Alpha defensin expression defines bone marrow derived T cells

Having characterized T cell pathways in an *in vitro* setting, we next aimed to understand how pathways are regulated when T cells migrate into tissue. For this, we analyzed a dataset from Szabo *et al*. (2019) whereby single cell sequencing was performed on unstimulated or anti-CD3 + anti-CD28 stimulated CD3^+^ T cells sorted from blood, bone marrow, lymph node, or lung of healthy donors. We identified 15 T cell types across the four tissue sites, with all annotated cells types and their markers shown in Figures 5A-B (Supplementary Figure 7). Given the scalability of SCPA, we then aimed to systematically characterize pathway activity in all T cell populations across all tissues and stimulation conditions to identify novel features of tissue derived T cells. For this, we used ∼3000 pathways taken from the Hallmark, KEGG, Reactome, Biocarta, PID, and Wikipathways database, comparing T cells from each tissue site to equivalent populations found in the blood, either in unstimulated or stimulated conditions. For example, unstimulated CD8^+^ Tem cells from the blood were compared to unstimulated CD8^+^ Tem cells from the lung, lymph node, and bone marrow, and so on (Figure 5C).

**Figure 5.**
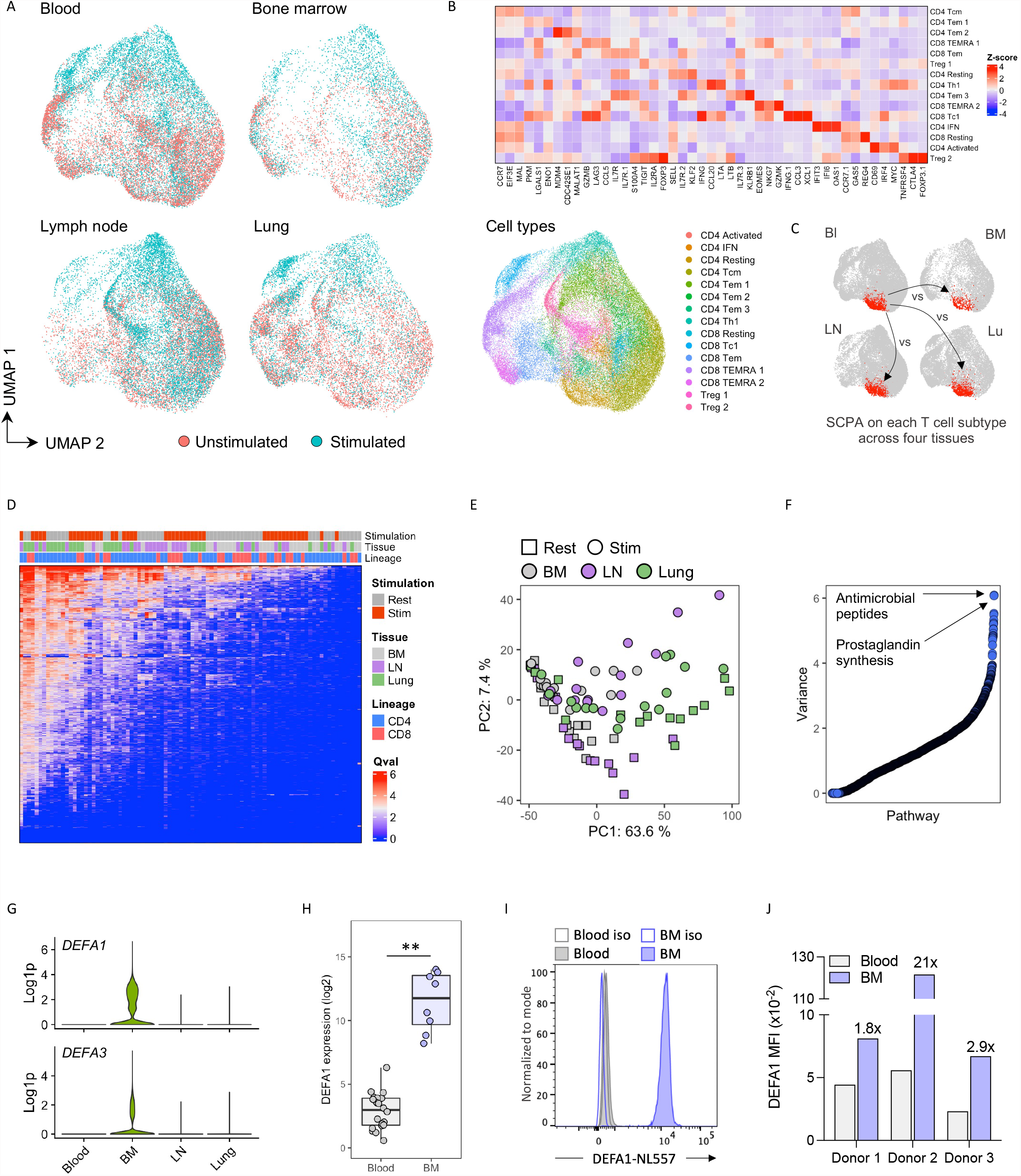
Alpha defensin expression defines bone marrow derived T cells (A) UMAP representations of stimulation effect and T cell subtypes across the indicated tissues. (B) Heatmap representation of markers from each cell subtype indicated in the UMAP plots (C) Outline of SCPA comparisons of cell types between tissues. Each cell type in each tissue was compared to the equivalent population found in the blood, both under resting and activated conditions. (D) Heatmap of Qvals generated by SCPA after pathway comparisons outlined in (C), with tiles above the heatmap representing selected metadata features of each column. (E) PCA of Qvals generated in (D), highlighting the effect of stimulation and tissue location of each cell type. (F) Dot plot showing the variance of all pathways across tissue sites and stimulation conditions (G) DEFA1 and DEFA3 expression in all T cell subtypes grouped together, split by tissue site. (H) DEFA1 expression in CD4^+^CD45RO^+^ T cells derived from blood or bone marrow, acquired from dataset GSE50677 (I-J) Flow cytometry analysis of DEFA1 expression in CD3^+^ T cells from bone marrow compared to blood derived CD3^+^ T Cells from age and sex matched healthy donors. Iso = isotype control. Data were acquired across three independent experiments

We were therefore able to characterize the changes in pathway activity when T cells are present in each tissue, as well as how they respond to stimulation. Globally, we observed that T cells present in the lung show the largest alteration in pathway activity compared to those from the lymph node and bone marrow (Figure 5D). Interestingly, a large source of variance within the data originated from the tissue site T cells were derived from, in addition to stimulation, suggesting that tissue migration imprints significant altered transcriptional profiles on cells even prior to activation (Figure 5E).

Having first mapped the global pathway signatures of tissue-derived T cells, we next aimed to identify pathways that were specific to T cells residing within the distinct tissue sites. To do this, we calculated the variance in the SCPA Q-values of each pathway across tissues and cell type (variance of rows in Figure 5D heatmap) to identify the pathways with the most variable effect sizes across the T-cell populations being compared. One of the pathways showing the largest perturbation were those involved in translation and ribosome assembly, with cells from the lung showing a broad downregulation of ribosomal subunit genes when compared to cells from other tissue sites (Supplementary Figure 7B). Furthermore, T cells from the tissue also showed high variance in arachidonic acid-derived prostaglandin synthesis and regulation, largely driven by expression of annexin family genes (*ANXA1-2*) that inhibit PLA2 activity, but also *S100A10* and *S100A6* that regulate annexin activity (Supplementary Figure 7C). However, the pathway with the most variable effect size was, unexpectedly, involved in antimicrobial peptide production. In assessing expression of genes from this pathway, this signature was driven by alpha defensin molecules DEFA1 and DEFA3, whose expression is currently thought to be limited mainly to neutrophils and mucosal epithelia. Moreover, assessing tissue specificity, this signature was exclusive to bone marrow-derived T cells, suggesting a tightly controlled tissue specific feature of T cells. To further validate our finding, we analyzed a previously published microarray dataset from Okhrimenko et al. (2014) of T cells sorted from the bone marrow and blood of healthy donors. In agreement with our data, we saw significantly increased transcript expression of DEFA1/3 in T cells from the bone marrow when compared to the blood (Figure 5H). Furthermore, to confirm this signature at the protein level, we sourced bone marrow of healthy human donors and compared DEFA1 expression to blood derived T cells from age and sex matched controls. DEFA1 was expressed almost exclusively in CD3^+^ T cells from the bone marrow when compared to CD3^+^ T cells from the blood (Figures 5I-J). These data therefore uncover a novel feature of bone marrow derived T cells; namely the expression of alpha defensin molecules.

In sum, we generate a global map of pathway activity in tissue derived T cells, identify a broad perturbation of pathway activity in T cells from the tissue, especially in lung derived T cells, and further discover a previously unrecognized alpha defensin expression profile in T cells from the bone marrow.

## Discussion

Here we introduced SCPA as a novel and effective approach to pathway analysis in scRNA-seq data. We utilized SCPA to systematically map pathway activity in T cells across time, between T cell populations, and across tissues, uncovering novel regulatory mechanisms of T cells.

Our approach to pathway analysis described in this work harnesses a statistical framework that considers changes in the multivariate distribution of all the genes of a given pathway as the primary statistic for judging biological relevance. This approach is highly robust, and does not depend on parametric assumptions on the gene expression distribution. By design, it is sensitive to distribution changes (Mukherjee *et al*., 2020) that would otherwise be missed using enrichment profiles. As an illustration, when considering a situation in comparing healthy and disease samples, whereby a pathway shows significant changes in its multivariate distribution but lacks enrichment of pathway genes in any given sample; this would still indicate substantial pathway perturbation. However, given the lack of mean differences, this would not be identified using current enrichment methods. We highlighted this unique feature of SCPA when we identified the requirement of cell-intrinsic arachidonic acid metabolism for T cell activation and effector function, which was not considered significant by GSEA. Of note, we found that many additional pathways demonstrated similar profiles (e.g. Figure 4D), which provides candidate pathways for future work. Further still, even though our statistical framework assesses distribution changes, gene sets that do show enrichment also necessarily show alterations in multivariate distribution, meaning that SCPA captures both aspects of pathway activity. This is also likely the reason why SCPA was able to identify a greater number of significant viral pathways when compared to other pathway analysis tools in our benchmarking (Figure 1). We therefore argue that assessing changes in multivariate distribution more accurately reflects what should be considered interesting when addressing pathway level gene signatures.

Employing SCPA to our scRNA-seq resource provided a systems level view of T cell transcriptional regulation upon activation, highlighting a number of interesting signatures. In contrast to current approaches that are largely restricted to two-sample comparisons, SCPA can assess gene set distributions across a multisample input simultaneously and is well suited to addressing experimental designs with greater than two conditions. Here we used SCPA over such a multisample design, assessing three early time points across T cell activation. We identified a core set of pathways that are shared upon T cell activation across naïve and memory CD4^+^ and CD8^+^ T cells. This led to the discovery of type I IFN signaling as a central transcriptional module in CD4^+^ T cell populations. T cells are responsive to exogenous IFNα (Huber and Farrar, 2011) but are not known to produce IFNα themselves. As the design of our T cell activation experiments excluded an exogenous source of IFNα, we concluded that the type I IFN signature must be rooted in an autocrine mechanism. Although limited in scope, our functional experiments supported such an idea as proof-of-principle: we demonstrated IFNα secretion upon T cell activation and a dependency on intrinsic IFNα for survival, at least *in vitro*. Indeed, we had previously established such a concept for another cytokine that was thought not to be produced by T cells; namely IL-1β. T cells produce IL-1β in very low amounts, yet intrinsic IL-1β controls the magnitude of Th1 responses (Arbore et al., 2016) and exogenous IL-1β cannot compensate for this function. Though IFNα is known to aid T cell survival (Marrack *et al*., 1999), a role for T cell-autocrine type I IFN may have previously been overlooked, as human T cells produce also only small quantities of this cytokine (Figure 3I). Additionally, *Ifna*^*–/–*^ mice do not exist, with IFNα targeted studies instead utilizing *Ifnar1*^*–/–*^ animals, or depleting classical sources of IFNα, such as pDCs, which makes it unsuitable to pinpoint T cell-autocrine IFNα functions. Two additional observations we made with regards to IFN biology will be worthy to investigate further. First, our data indicated that T cells produce IFNA with at least some specificity toward IFN21 – though our primers also showed an overlap with IFNA1/13 – and not other IFNA genes including IFNA4, IFNA7 or IFNA17. This is interesting in view the of emerging realization that different type I IFN subtypes can have diverse effects on CD8^+^ T cell antiviral responses (Dickow et al., 2019). Furthermore, we showed that while all T cells harbor or acquire a strong IFN pathway signature, the respective gene transcription profiles induced are distinct among T cell subpopulations. We noted a specific demarcation between (memory) Tregs and non-regulatory T cells, suggesting tailored and selective responses downstream of the IFN receptor between lineages of inflammatory and regulatory T cells.

SCPA also highlighted metabolism as a pathway class with large distribution changes upon T cell activation. Although it is well established that all aspects of the T cell life cycle are controlled by metabolic events (Buck et al., 2015) most existing works focus on the analysis of one or a few select metabolic pathways in a given study. Here we provided a means for comprehensive pathway analyses of scRNA-seq data that can distinguish changes in the landscape of metabolic pathways upon cell activation, over time, or across stimulation conditions. Across naïve CD4^+^ T cell activation we confirmed that fundamental metabolic pathways, such as glycolysis and OXPHOS, featured prominently as expected. However, we also uncovered several metabolic pathways previously not associated with T cell metabolism. One such pathway was that of arachidonic acid metabolism, which we showed was necessary for proper T cell activation and IFNγ/IL-10 production. Though we did not define the exact mechanism of this regulation, downstream metabolites of arachidonic acid generated by immune cells, including as prostaglandins, prostacyclins, and leukotrienes, impact T cell function at several levels (Maseda et al., 2019). Much of the literature on arachidonic acid metabolite production focuses on innate immune cells, though our work here shows that T cells produce and utilize arachidonic acid via phospholipase A2 to induce proper T cell activation *in vitro*. In addition, we also observed upregulation of genes involved in the inhibition of PLA2 (ANXA and S100 family members) in T cells from the tissue, suggesting that inhibition of arachidonic acid metabolism *in vivo* could contribute towards confining T cell activation *in vivo*. Indeed, a recent paper has demonstrated that T cells from patients with rheumatoid arthritis are hyper-responsive to exogenous arachidonic acid, resulting in increased calcium flux and pERK signaling (Ye et al., 2021), hinting that dysregulation of this pathway in T cells could play a role in inflammatory disease.

In addition to mapping metabolic pathways across T cell activation, we compared metabolic transcription across quiescent T cell subpopulations and demonstrated that memory CD4^+^ T cell lineages maintain different levels of metabolic poise. We found that whilst Tcm cells display a broadly similar metabolic profile relative to naïve T cells, quiescent Th1 and Th2 cells maintain altered levels of metabolic transcription across a broad range of metabolic pathways. Whilst previous work has shown that bulk memory T cells exhibit higher levels of glycolysis and OXPHOS relative to naïve cells (Gubser *et al*., 2013), our findings here suggests that this heightened metabolic activity may be predominantly driven by a minority of memory CD4^+^ T cell subpopulations, rather than memory T cells as a whole. Further, we suggest that the metabolic distinction between naïve and memory T cells may be larger than previously appreciated, noting that a broad range of metabolic pathways outside of glycolysis and OXPHOS were differentially regulated. One such observation was that a number of glycogen-related metabolic pathways were active in CD4^+^ effector memory populations relative to Tcm cells. The generation of glycogen stores in dendritic cells supports their early effector functions via contributions to glycolytic reprogramming and mitochondrial respiration after toll-like receptor ligation (Thwe et al., 2017). Further, glycogen breakdown has been shown to support CD8^+^ T cell memory homeostasis and survival (Ma et al., 2018). Although speculative at this point, the glycogen signature in effector memory T cells could represent an important component of the heightened metabolic poise that enables these cells to respond rapidly to stimulation. Overall, our work here focused on T cells, though we anticipate similar systematic approaches to immune metabolism using scRNA-seq will reveal important and previously undiscovered aspects of immune regulation across other cell types.

In addition to our *in vitro* work, we performed a systems level analysis of pathways in *ex vivo* T cells derived from different tissue sites. This was possible given that SCPA provides a computationally scalable approach to identify pathway signatures in vast amounts of multisample transcriptomic data. In this, we analyzed around 3000 pathways in 15 cell types across multiple tissues and stimulation conditions. Surprisingly – as it is currently thought that defensin production is mainly restricted to innate immune cells and epithelial cells (Xu and Lu, 2020) – we identified alpha defensin expression, through DEFA1 and DEFA3, as a unique feature of T cells from the bone marrow. To our knowledge, there is no precedent for alpha defensin expression by αβT cells. Previous work has been unable to detect alpha defensin expression in CD8^+^ T cells from the blood by mass spectrometry (Mackewicz et al., 2003), with similar results in two αβ T cell lines (Agerberth et al., 2000), but did detect alpha defensin in CD3^+^ T cells, likely coming from the γθ fraction. These findings are in line with our data, suggesting that alpha defensin expression seems to only be upregulated in αβ T cells after migration into the bone marrow, and not in other tissues. We did not uncover the reason for alpha defensin expression in αβ T cells, however, the antimicrobial nature of defensins possibly contributes to general sterility of the bone marrow microenvironment. Nonetheless, the immune regulatory properties of defensins are broad, meaning defensin expression could influence a multitude of processes such as cell migration, cytokine release, and proliferation (Fruitwala et al., 2019; Xu and Lu, 2020), and future work defining the precise role of T cell derived alpha defensins is required.

In summary, by developing and employing SCPA, we uncovered novel cytokine and metabolic pathways engaged during early T cell stimulation and provided comprehensive global signatures of pathway alterations among distinct T cell subpopulations. Overall, this work outlines the power of systematic approaches to uncover novel regulatory pathways across a wide range of cell types and tissues. Furthermore, though our work here focused on T cells from healthy donors, this approach will be invaluable in characterizing pathway perturbations in a disease setting, generating an unbiased analysis of the relative cell specific dysfunctions contributing to disease pathogenesis, alongside providing therapeutic targets.

## Methods

### Human CD4^+^ and CD8^+^ T cell Purification and In Vitro Activation

Human bulk CD4^+^ or CD8^+^ T cells were isolated from PBMCs, obtained from freshly drawn blood after centrifugation using Lymphoprep separation medium (Corning, Vienna, VA) using either the MACS Human CD8^+^ T cell Isolation Kit (130-096-495) from Miltenyi Biotech, (Bergisch Gladbach, Germany), the Negative Selection EasySep CD4^+^ T cell kit (17951) or RosetteSep Human CD4^+^ Cell Isolation Kit (15062) from Stemcell Technologies (Vancouver, Canada) according to the manufacturers’ instructions. Enriched purified CD4^+^ or CD8^+^ T cells were then stained with a combination of antibodies to CD45RA-FITC, CD45RO-, CD4-PE, CD8-PercpCy5.5, CD56-APC for 30 min at 4°C and subsequently sorted into naive or memory CD4^+^ or CD8^+^ T cell subpopulations using the Cell Sorter SH800S (Sony Biotechnology, Inc., San Jose, CA). Cell purity was consistently > 99 %. Purified naive or memory CD4^+^ or CD8^+^ T cells were activated for indicated time points in 48-well culture plates (Greiner, Monroe, NC) at 2.5 – 3.0 × 10^5^ cells/well in media containing 25 U/ml recombinant human IL-2 in an incubator at 37°C and 5 % CO2. Plate bound anti-CD3 and anti-CD28 were used to stimulate CD4^+^ (2µg/ml both), and CD8^+^ (0.25 and 2 µg/ml, respectively) T cells.

### Bone marrow samples

Human bone marrow aspirates were obtained from healthy volunteers after informed consent in accordance with the Declaration of Helsinki, under an Institutional Review Board-approved clinical protocol (NCT00442195). Mononuclear cells were separated using Ficoll-Hypaque density gradient centrifugation (MP Biomedicals) and stored in CryoStor CS5 freezing medium (Biolife Solutions) under liquid nitrogen vapor phase until use. For analysis, cells were thawed in PBS supplemented with 2mM EDTA, 0.5% HSA (Baxter Healthcare Corporation), 10 units/mL DNase (Genentech, Inc.) and 2.5mM MgCl_2_.

### Mice

Both wild-type and IFNAR1^-/-^ mice were on a C57BL/6 background, with IFNAR1^-/-^ mice previously described in (REF). Splenic single cell suspensions were generated, and red blood cells lysed using ACK lysis buffer (Life Technologies). CD4^+^ T cells were isolated by negative selection using the Stem Cell Technologies EasySep Mouse CD4^+^ T Cell Isolation Kit (Tukwila, WA), resulting in a CD4^+^ purity of 95-98%, and a CD3^+^ purity of ∼99%. CD4^+^ T cells were then plated at 180,000 cells per well in a 96-well plate and activated with plate-bound anti-CD3 (2µg/ml) and soluble anti-CD28 (1µg/ml) for the indicated time before harvesting and staining with near-IR fluorescent reactive dye (Invitrogen).

### Flow cytometry and cell sorting

Cells were harvested after stimulation and viability of cells post activation was measured using the LIVE/DEAD™ Fixable Aqua Dead Cell Stain Kit (Thermo Fisher Scientific). Surface staining of antigen was performed at 4°C for 20 minutes. For intracellular antigens, cells were fixed and permeabilized using the FOXP3/Transcription factor staining kit (eBioscience), and incubated with the appropriate antibody for 1 hour at 4°C. Acquisition of cells was performed using the BD FACSCantoTM II Flow Cytometer and subsequent analysis was performed using FlowJo (v9 or v10). The clone and dilution of antibodies used for flow cytometry were as follows: CD4-Pacific Blue (Biolegend, OKT4, 1:200), CD69-APC (Biolegend, FN50, 1:200), CD45RA-APC (Biolegend, HI100, 1:200), CD45RO-FITC (Biolegend, UCHL1, 1:200), DEFA1 (R&D Systems, 2µg/ml), anti-sheep IgG-NL557 (R&D Systems, 1:500). IFNγ expression was analyzed using the IFN-γ Secretion Assay (PE, Miltenyi Biotec) following manufacturers instruction.

### Library sequencing and preprocessing steps for the single cell transcriptomics data

We prepared single cell transcriptome libraries using the 10xGenomics Chromium Single Cell v3 kit, in accordance with the company’s manual. Subsequently, the single cell libraries were sequenced by the NovaSeq 6000 sequencing system (Illumina, Inc., San Diego, CA, USA) with reads per cell targeted to be at a mean of 50,000 reads for each sample sequenced. Initial quality control was performed on the resulting scRNAseq data from each sample using Cell Ranger (Version 2.1.0). All data processing parameters and scripts used for processing all samples can be found at https://github.com/jackbibby1/scpa_paper. Briefly, we excluded any cells with > 10% of reads mapping to mitochondrial genes, and filtered cells based on number of genes expressed. Filtered count matrices were then normalized by total UMI counts, multiplied by 10,000 and transformed to natural log space. The top 2000 variable features were determined based on the variance stabilizing transformation function (FindVariableFeatures) by Seurat with default parameters. Any cycle genes were removed from the variable features list to reduce cell-cycle associated variation, prior to downstream analysis. For each sorted T cell population, samples were integrated across time points using canonical correlation analysis (CCA), resulting in four integrated datasets. For example, naïve CD4^+^ T cells at 0, 12, and 24 hours were integrated, followed by memory CD4^+^ T cells, and so on for CD8^+^ T cells. Variants arising from cell cycle phases and the percentage of mitochondrial genes were regressed using by the ScaleData function in Seurat. Principal component analysis (PCA) was then performed, and the top 30 Principal components (PCs) were included in a Uniform Manifold Approximation and Projection (UMAP) dimensionality reduction. Clusters were identified by canonical marker genes on a shared nearest neighbor (SNN) modularity graph using the top 30 PCs and the original Louvain algorithm. Markers for each population were calculated using the FindAllMarkers() function within Seurat, with parameters: min.pct = 0.1, logfc.threshold = 0.25, only.pos = T. Definition of each cell population was done using literature associated markers, and all markers for each population can be found in Supplementary Table 1.

### Pathway analysis using SCPA

Scripts for replicating all figures in the paper can be found at https://github.com/jackbibby1/scpa_paper. For pathway analysis, log1p normalized data was extracted from each relevant population and pathways were pulled from the molecular signatures database using the msigdbr package within R. Comparisons were then done using the compare_pathways() function within SCPA using default parameters. Data processing and visualization was then performed using Seurat (Satija et al., 2015), ggplot2 (Wickham, 2016), ComplexHeatmap (Gu et al., 2016), and dyno (Saelens *et al*., 2019) R packages.

### Benchmarking pathway analysis methods

3 publicly available datasets were used for comparisons, under the accession numbers: GSE122031, GSE148729, and GSE156760 (containing 2 datasets used here). For Enrichr and DAVID, data were split between mock or virally infected cells, and differentially expressed genes were calculated using the FindMarkers() function within Seurat, using a logFC threshold of 0.4. Differentially expressed genes were then input into DAVID and Enrichr, and results were filtered to include GO Biological Process gene sets. For SCPA and GSEA, data were split by between mock and virally infected cells, cell numbers were downsampled to 500 cells per sample, and then expression files were directly used downstream. GSEA was ran using default parameters: ‘phenotype’ permutation, ‘weighted’ enrichment statistic, Signal2Noise ranking metric, and gene sets with < 15 genes or > 500 genes were excluded. SCPA was ran using default parameters, excluding gene sets with < 15 genes or > 500 genes. Viral gene sets were defined by any pathways containing the string “VIR” in their pathway name, within the GO Biological Process gene set list, and all viral pathways can be found in Supplementary Table 4.

### Trajectory inference analysis and pseudotime SCPA

All scripts for the trajectory inference analysis can be found at https://github.com/jackbibby1/scpa_paper. The R package dyno was used to calculate trajectories and discrete nodes across pseudotime. Briefly, the top 1000 most variable genes were taken from naïve T cells across 0, 12, and 24 hours (after excluding regulatory T cells) and used in the downstream analysis. The naïve CD4^+^ T cell trajectory was then calculated using the infer_trajectory() function within dyno, using the ti_slingshot() method. Milestones along the trajectory were calculated using the group_onto_nearest_milestones function within dyno. After assignment, cells were then extracted based on their milestone identifier, and these discrete populations were subsequently used for input into SCPA. A list of manually curated 243 metabolic pathways were used for the pseudotime pathway analysis, which can be found in Supplementary Table 3.

### Graph-based test used in SCPA

Given K multivariate probability distributions F_1_, F_2_, …, F_K_, the K-sample problem is to test the hypotheses

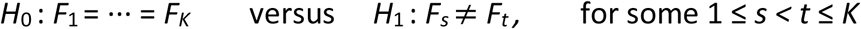

based on collections of independent observations – *χ*_1,_ *χ*_2,_ …, *χ*_*K*_– coming from F_1_,F_2_,…,F_K_ respectively. In the single cell setting, F’s correspond to the tissue samples, and *χ*’s denote the number of cells sequenced for a given condition K. For example, in this work we are interested in comparing the multivariate distribution of a gene set (corresponding to a particular pathway) across K=3 conditions, F_1_ (0h), F_2_ (12h) and F_3_ (24h), for a given cell type.

Several tests exist for nonparametric testing in the 2-sample framework that are based on the idea of constructing graphs on the pooled sample *χ*_1_ ∪ *χ*_2_ ∪ … ∪ *χ*_*K*_, such as nearest neighbor graphs (Henze, 1988; Schilling, 1986), minimal spanning trees (Friedman and Rafsky, 1979) or graphs based on optimal non-bipartite matching (Rosenbaum, 2005). Inspired by the latter method and ideas developed in another recent paper (Chen and Friedman, 2017), we developed a universally consistent and distribution-free multisample test, i.e. the null distribution of the test statistic does not depend on the distribution of the data (Mukherjee *et al*., 2020). Briefly, our novel multisample crossmatch test, based on the Mahalanobis disparity, begins by pooling the samples *χ*_1,_ *χ*_2,_ …, *χ*_*K*_ corresponding to the distributions F_1_,F_2_,…,F_K_ (respectively) together. We then construct a minimum non-bipartite matching graph on the pooled sample, i.e., a matching on {*χ*_1_ ∪ *χ*_2_ ∪ … ∪ *χ*_*K*_}, which minimizes the sum of the Euclidean lengths of all the edges. For each 1 ≤ s < t ≤ K, the (s,t)-cross count (*a*_*st*_), is then defined as the number of matched edges in the pooled sample {*χ*_1_ ∪ *χ*_2_ ∪ … ∪ *χ*_*K*_} with one endpoint in *χ*_s_ and the other endpoint in *χ*_t_. The test statistic is then defined as:

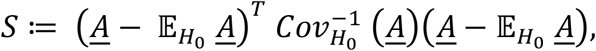

where 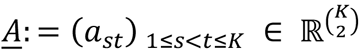 denotes the vector of cross-counts, and 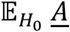 and 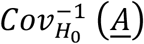 denote the null mean vector and the null covariance matrix of *<u>A</u>*, respectively.

*S* is distribution free under H0, and exact expressions for the entries of *A* can be computed from a given dataset. Instead of using the exact null distribution of *S* its weak limit, the chi-squared distribution with 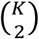 degrees of freedom can be used for computing the test cut-off. Since *S* is a measure of distance between 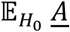 and *<u>A</u>*, the null hypothesis is rejected when *S* is “sufficiently large”, i.e. exceeds a certain null hypothesis p-value cut-off. Additional theoretical details are described and proved in the associated statistical work {Mukherjee, 2020 #6}.

The *S* test statistic above is used to construct the *Q* value used in this paper and in scpa for ranking of pathways by degree of distributional change. First, the p-value *p* of *S* is computed as

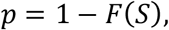

where *F* is the *χ*^2^ distribution with 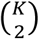 degrees of freedom. The *Q* value is then the transformation

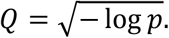

### qRT-PCR

CD4^+^ T cells were either left unstimulated or stimulated for 12 or 24 hours with antiCD3 + anti-CD28 (both 2µg/ml). Cells were harvested, washed in PBS, lysed in TRIzol (Ambion), and stored at −80°C until used. RNA was extracted using the Direct-zol RNA MiniPrep kit (Zymo research), and subsequent reverse transcription was performed with 200ng of RNA using the High-Capacity RNA-to-cDNA kit (applied biosystems). qPCR analysis was then performed using the TaqMan Universal Master Mix II, no UNG (applied biosystems), with FAM labelled primers for IFNA1/13/21 (Hs04190680_gH), IFNA4 (Hs01681284_sH), IFNA7 (Hs01652729_s1), IFNA17 (Hs00819693_sH), and a VIC labelled primer for RPL7 (Hs02596927_g1) following manufacturer’s instructions.

### CRISPR-Cas9 deletion of IFNα

crRNA selection: Due to potential for gene redundancy, we designed a total of four crRNA targeting four *IFNA* genes using the Benchling (www.benchling.com) online platform (See Table 1). In order to target as many *IFNA* genes as possible, we considered using crRNA with a relatively high on-target potency against other *IFNA* genes, explaining the low off-target score. crRNA suspected to have off-target effects directed against genes other than *IFNA* family members were excluded. As a result, we targeted 4 main IFNA genes, but this includes overlap with all IFNA genes apart from IFNA8. crRNAs were ordered from Integrated DNA Technologies (www.idtdna.com/CRISPR-Cas9) in their proprietary Alt-R format.

**Table1.**
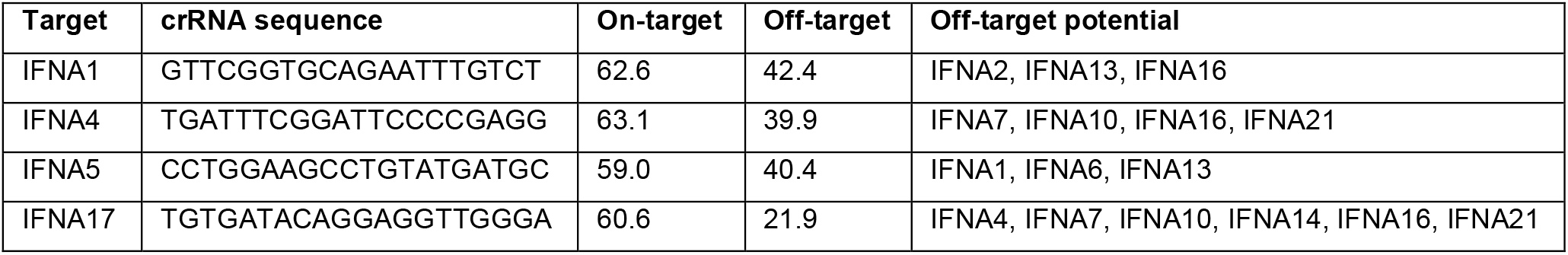
crRNA design including on and off target scores generated by Benchling

Preparation of cells: Human CD4^+^ cells were isolated from buffy coats using EasySep™ Human CD4^+^ T Cell Isolation Kit (Stemcell Tech, # 17952), resuspended with complete RPMI (cRPMI) media (supplemented with P/S, L-Glu, 10% FCS) and kept on ice until needed.

Preparation of crRNA–tracrRNA duplex: Each crRNA and tracrRNA (IDT, #1072534) were reconstituted at a final concentration of 160µM using nuclease-free duplex buffer (IDT, #1072570). For one electroporation, 1µL of crRNA was mixed to 1µL of tracrRNA in a sterile, RNAse-free PCR tube. Oligos were annealed at 95°C for 5 min in PCR thermocycler and slowly cooled down to 4°C. Precomplexing of Cas9/RNP: In the same PCR tube, 1.2µL of Cas9 nuclease (IDT, #1081059) was added, and the mixture was incubated 15 min at 37°C to form RNP complexes. The four RNP complexes against IFNA1, IFNA4, IFNA5 and IFNA17 were pooled and further kept on ice until needed.

Nucleofection: Up to 10 × 10^6^ of freshly isolated human CD4^+^ T cells were used per electroporation. Cells were transferred in 1.5 mL Eppendorf tubes and washed twice with PBS 1X (Gibco) to remove all trace of FCS. Immediately before electroporation, cells were resuspended with 10µL of primary cell nucleofection solution (P3 primary Cell 4D-Nucleofector X kit S (Lonza, # V4XP-3032), and mixed with RNP complexes. The mixture cells/RNP complex was then transferred into electroporation strip. Within 5 minutes, cells were electroporated using “EH-100” pulse on 4D-nucleofectore core unit (Lonza). Immediately after electroporation, 100 µL of pre-heated cRPMI + 20 UI/mL IL-2 was carefully added directly into each well of the nucleofection strip, and the strip was placed in tissue culture incubator for 15-30 min to allow for cell recovery. Cells were then transferred into a culture plate, resuspended at 2 × 10^6^ / mL in cRPMI + 20 UI/mL IL-2 and incubated at 37°C for 3 days before stimulation.

### T7 endonuclease assay

DNA extraction: DNA was extracted from control-RNP and IFNA-targeted cells using Quick-DNATM Microprep kit (Zymo Research, # D3021), and DNA concentration was calculated on Nanodrop OneC (Thermofisher).

PCR amplification: PCR primers were designed on Primer designing tool (NCBI platform) and purchased from IDT. Primer sequences were as follows: fwd: TGATCTCCCTGAGACCCACA rev: CAGGGGTGAGAGTCTTTGAAATG. PCR amplification was performed using the Q5® Hot Start High-Fidelity 2X Master Mix (NEB, M0494) following the protocol provided by the manufacturer. The hybridization temperature for each forward/reverse couple was calculated using the NEB Tm Calculator. PCR amplification products were later purified using DNA Clean & ConcentratorTM −5 (Zymo Research, # D4004) following the protocol provided by the manufacturer.

T7 endonuclease I digestion: 200 ng of purified DNA amplicons were annealed with 1X NEBuffer 2 (NEB, # B7002S) and total volume was brought up to 19 uL. Hybridization step was performed following manufacturer’s protocol on thermocycler machine. Upon hybridization step completion, mismatch double strand DNA was detected by adding 1 uL of T7 endonuclease I in the mixture and incubated 15 min at 37°C. The reaction was stopped by adding 1.5 uL of 0.25 M EDTA. The digested product was diluted 1:5 in molecular biology grade water and migrated on E-Gel™ EX Agarose Gels, 2% (Invitrogen, # G401002) for 10 minutes. Bands were revealed with ChemiDocTM Imaging System (BIO-RAD) using SYBR safe filter.

Gene modification was estimated using the following formula:

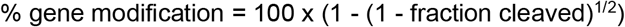

### Measurement of T cell supernatants

Supernatants from cultured T cells were harvested and stored at −20°C until analysis. Arachidonic acid measurement was done using the human arachidonic acid ELISA kit (Novus biologicals) following manufacturer’s instructions. Measurement of cytokines was done using the legendplex inflammation panel mix and match (IFNγ, TNFα, IL-10, Biolegend) or VeriKine-HS Human IFNα all subtype TCM ELISA (PLB assay science), following manufacturer’s instructions.

### OCR and ECAR Measurements

For analysis of OCR (in pMoles/min) and ECAR (in mpH/min), the Seahorse XF-96 metabolic extracellular flux analyzer was used (Seahorse Bioscience, North Billerica, MA). Naive or memory CD4^+^ or CD8^+^ T cells were resuspended in serum-free unbuffered RPMI-1640 medium (R1383; Sigma Aldrich) post activation and were plated onto Seahorse cell plates at 4 × 105 cells per well coated with Cell-Tak (CB-40241; Corning, Reinach, Switzerland) to enhance T cell attachment. Perturbation profiling of the use of metabolic pathways by T cells was achieved by the addition of oligomycin (O4876; 1 µM), Carbonyl cyanide-4-(trifluoromethoxy) phenylhydrazone (FCCP, C2920; 2 µM) and rotenone (R8875; 1 µM - all from Sigma Aldrich, St. Louis, MO). Metabolic parameters were then calculated based on the following formulas:

1. basal respiration =[OCR(basal-nc)] – [OCR(rotenone)]
2. ATP coupled respiration = [OCR(basal-nc)] – [OCR(oligomycin)]
3. maximal respiratory capacity = [OCR(peak-FCCP)] – [OCR(rotenone)]

## Data availability

All single cell data generated in the paper is available at [upload data]. We have included all raw files, and processed files containing count tables and annotation information for each sorted population, and processed R objects for each population. Single cell data using tissue derived T cells was taken from GSE126030. Microarray expression data from bone marrow T cells was taken from GSE50677. Datasets used in benchmarking was taken from GSE122031, GSE148729, and GSE156760.

## Code availability

Code used to generate all figures in the paper can be found at https://github.com/jackbibby1/scpa_paper. Tutorials and reference documentation for SCPA can be found at https://jackbibby1.github.io/SCPA

## Acknowledgements

This work was supported in part by the Intramural Research Program of the NIH, the National Heart, Lung, and Blood Institute (project number zia/hl006223 to C.K., Z99 HL999999 to A.L.), and 5R01-HG006137-07 and 1U2CCA233285-01 to N.R.Z. We thank Howard Young and Julio Valencia for providing the Ifnar1^-/-^ mice used in this study.

## Author Contributions

Cell culture, sorting, and preparation for single-cell analysis was performed by T.F., C.K., and E.E.W. Seahorse analysis was performed by N.K. and C.K. Bone marrow samples were provided by F.C and A.L. CRISPR experiments were designed by N.M. Analysis of single-cell data, SCPA package development, and other wet lab experiments were performed by J.A.B. The statistical framework underlying SCPA was developed by D.A., S.M., and N.Z. The manuscript was written by J.A.B., D.A., B.A., C.K., and N.Z., and the project was supervised by C.K. and N.Z.

---

**Supplementary Figure 1.**
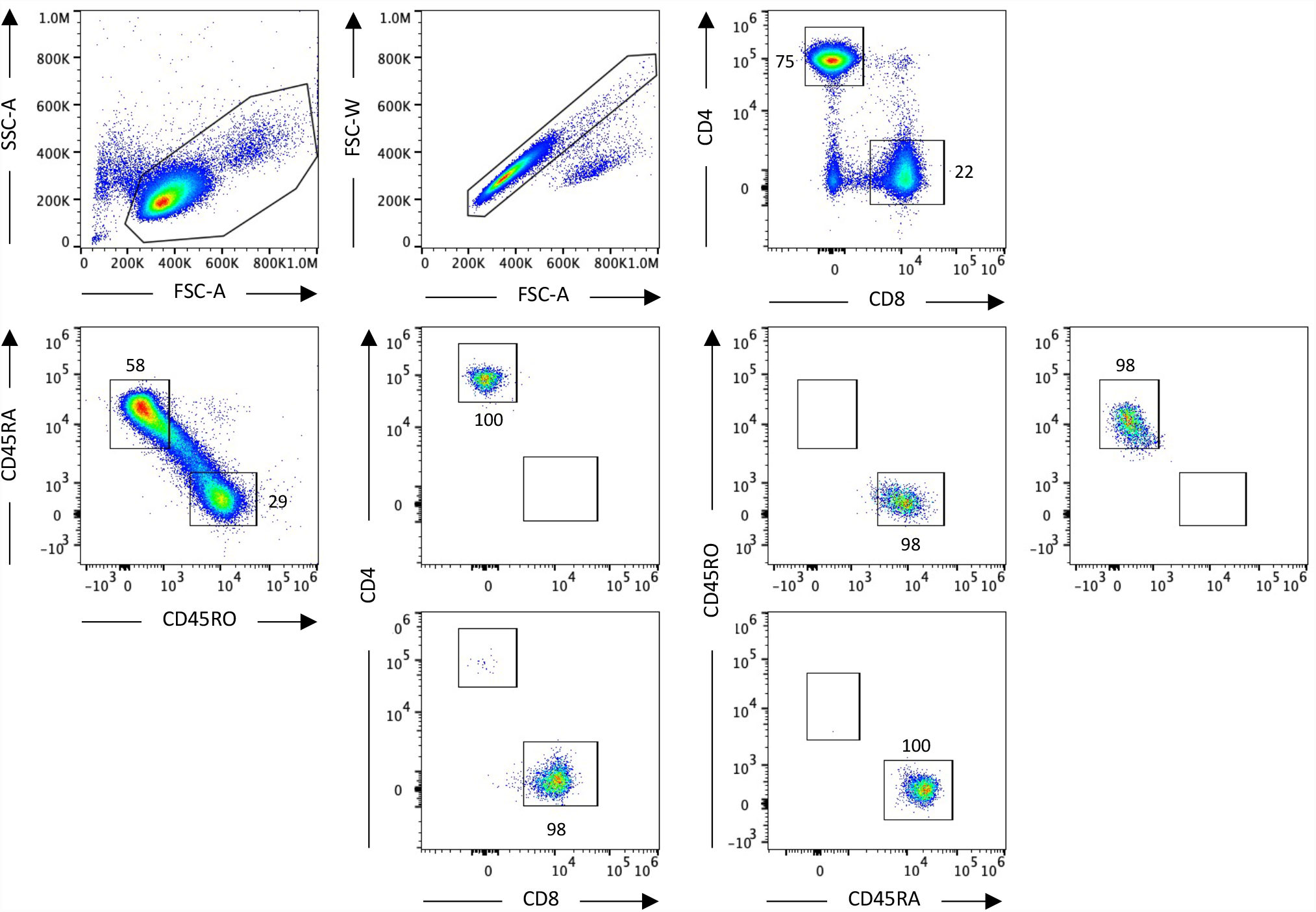
Flow cytometry gating and purity of T cell populations. T cells were gated as follows: FSC-A vs SSC-A for cells > FSC-W vs FSC-A for single cells > CD4^+^ vs CD8^+^ for T cell populations > CD45RA^+^ vs CD45RO^+^ for naïve and memory populations. Purity was then assessed post sort, showing high purity of T cell populations.

**Supplementary Figure 2.**
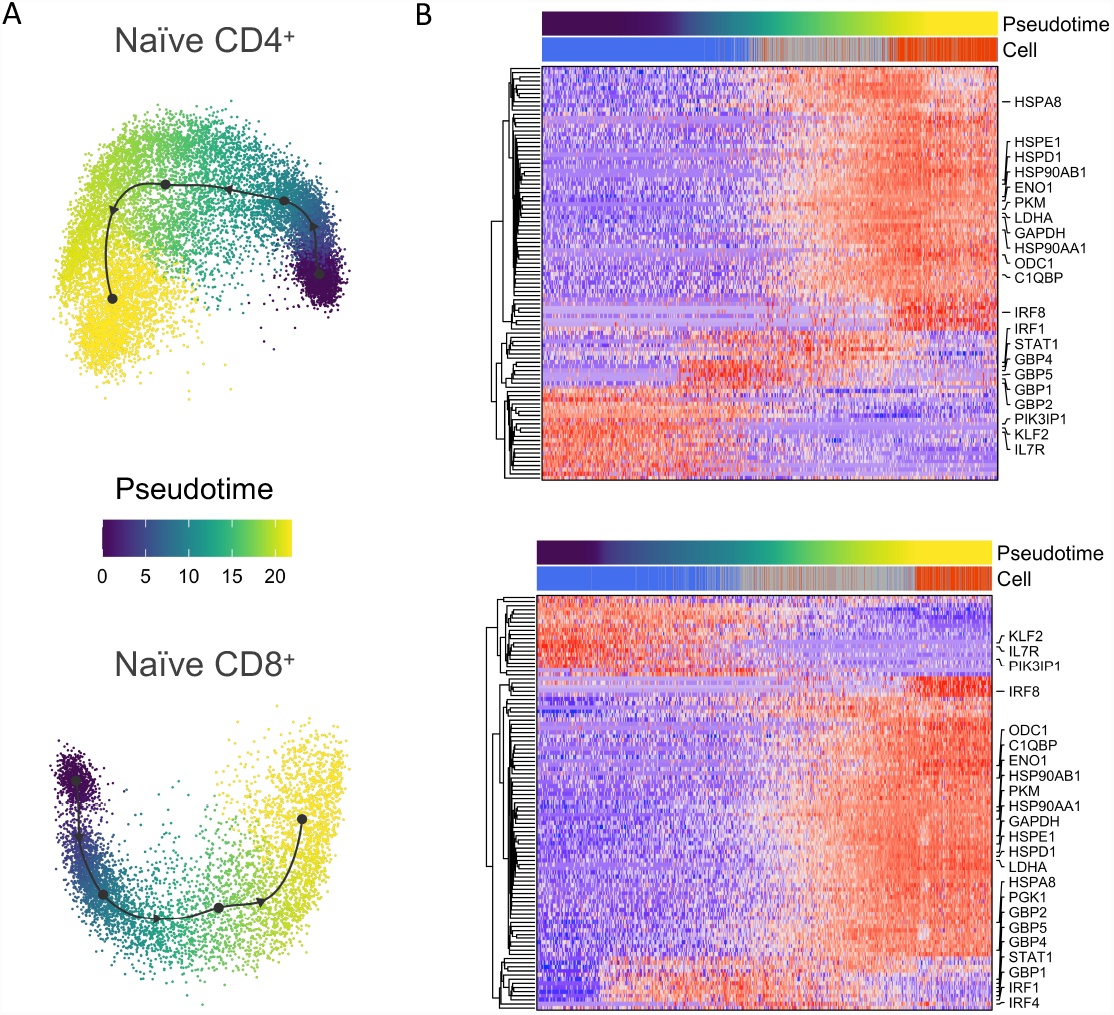
Pseudotime analysis of naïve CD4+ and CD8+ T cell activation (A) Pseudotime trajectory of naïve CD4^+^ and CD8^+^ T cells modelled using slingshot. (B) Heatmap representation of defining genes across pseudotime. Bars above the heatmap represent pseudotime value and cell identities taken from the UMAP: blue = resting, gray = intermediate, red = activated.

**Supplementary Figure 3.**
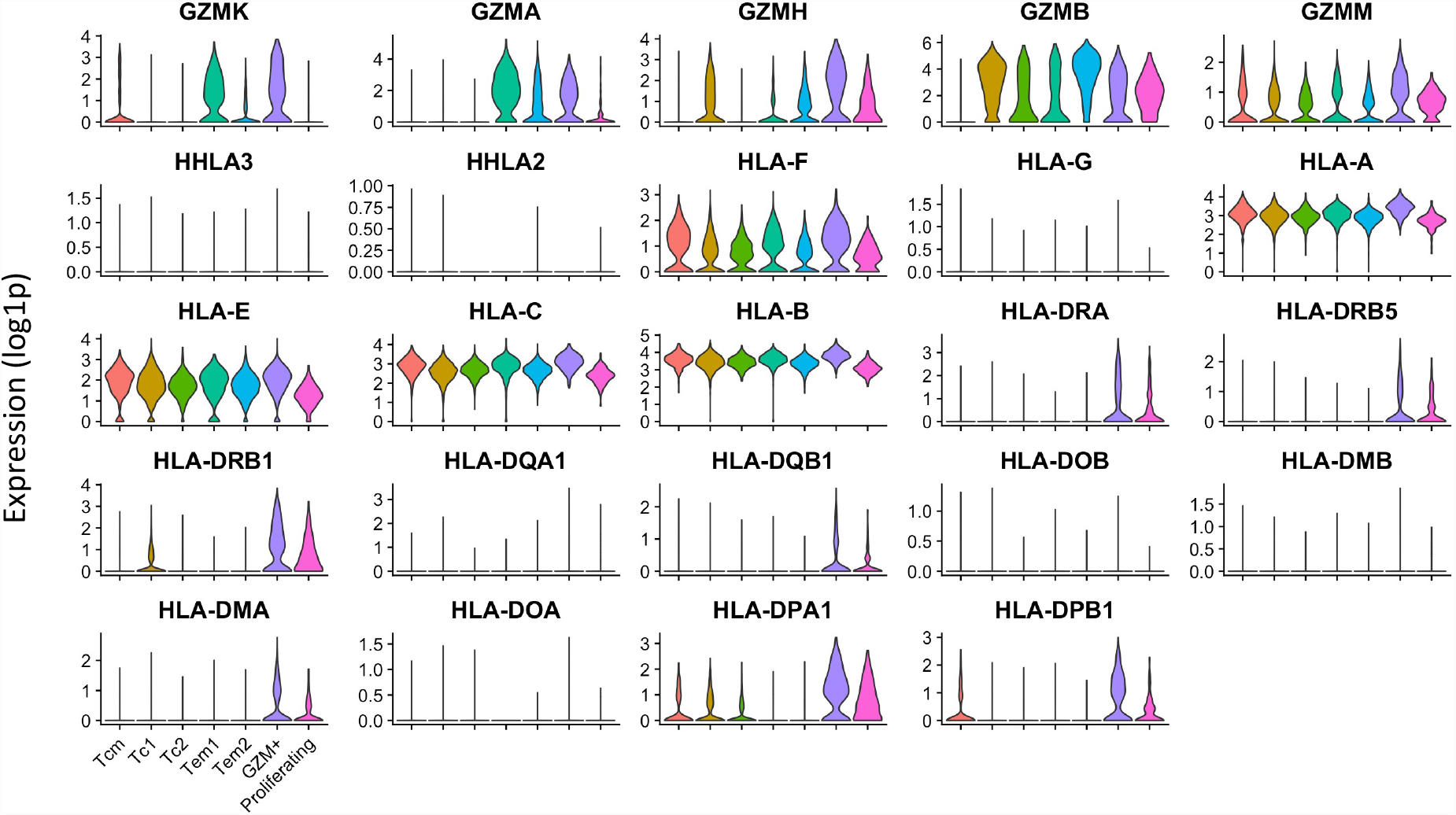
GZM and HLA expression in GZM+ CD8+ T cells Relative expression of GZM and HLA family member genes in memory CD8^+^ T cell populations

**Supplementary Figure 4.**
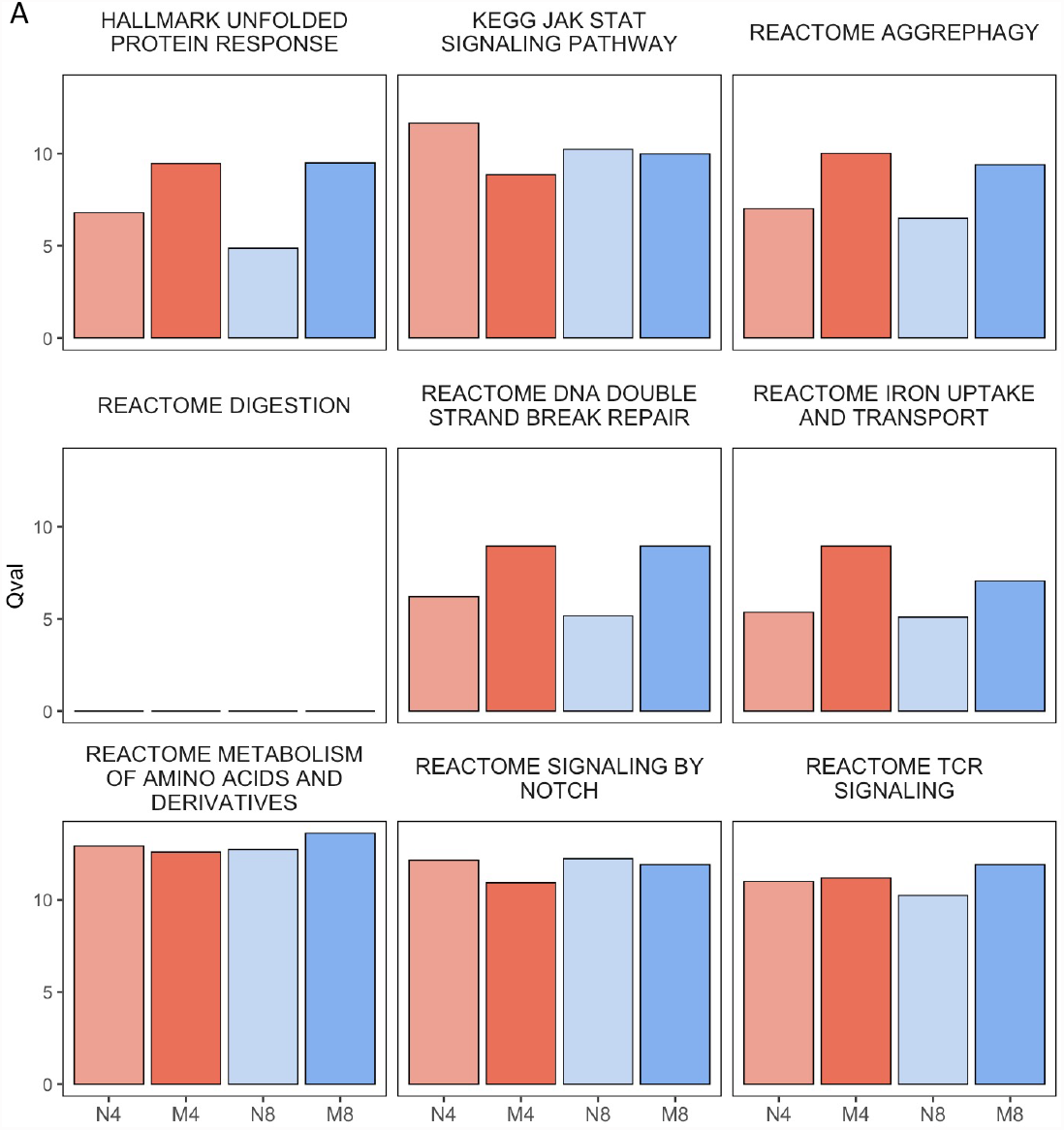
Global analysis of pathways upon T cell activation (A) Qvals for each pathway across T cell activation in naïve and memory CD4^+^ and CD8^+^ T cell populations. The four broad cell types were split across 0, 12, and 24 time points, and SCPA was used to compare pathway activity.

**Supplementary Figure 5.**
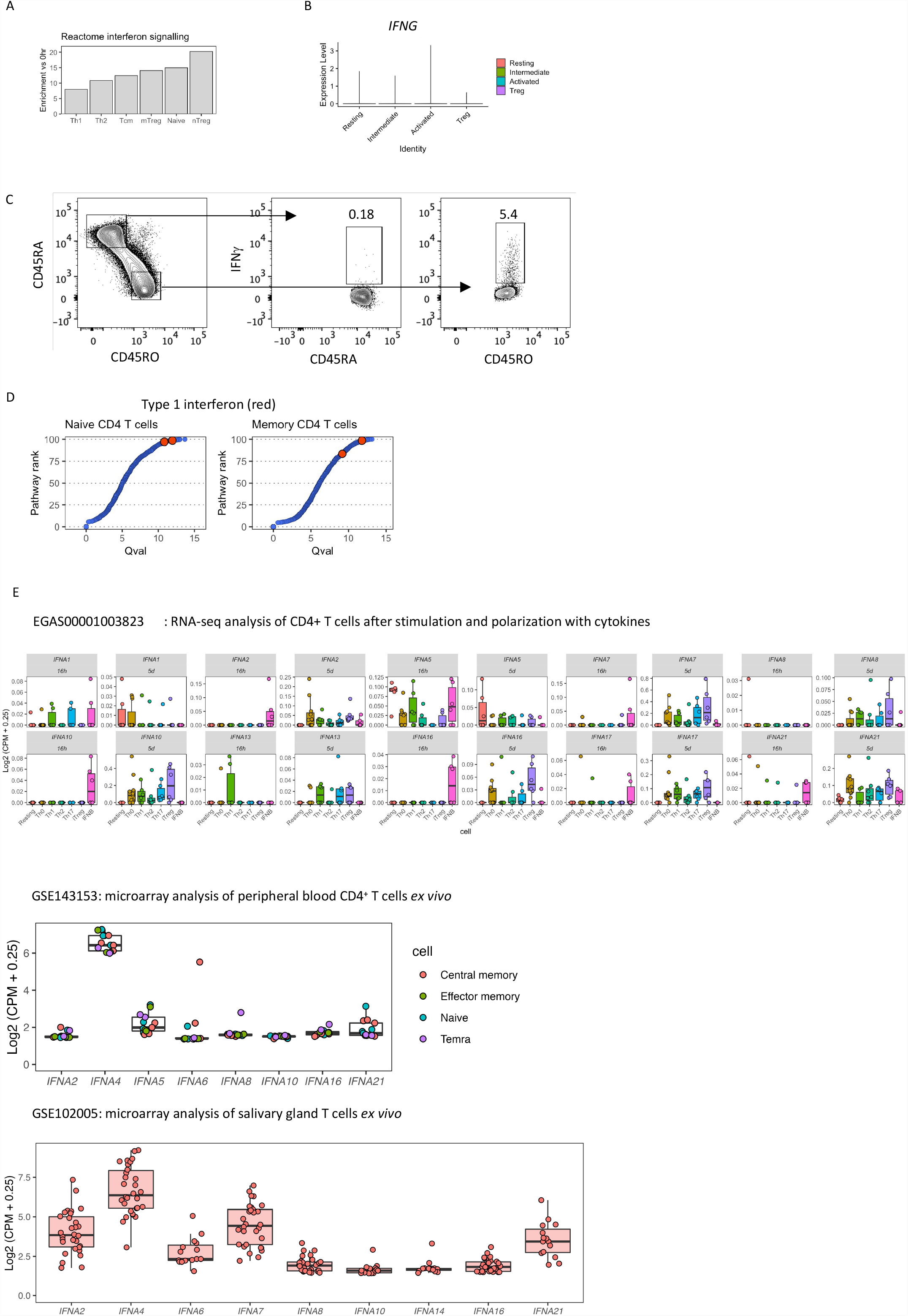
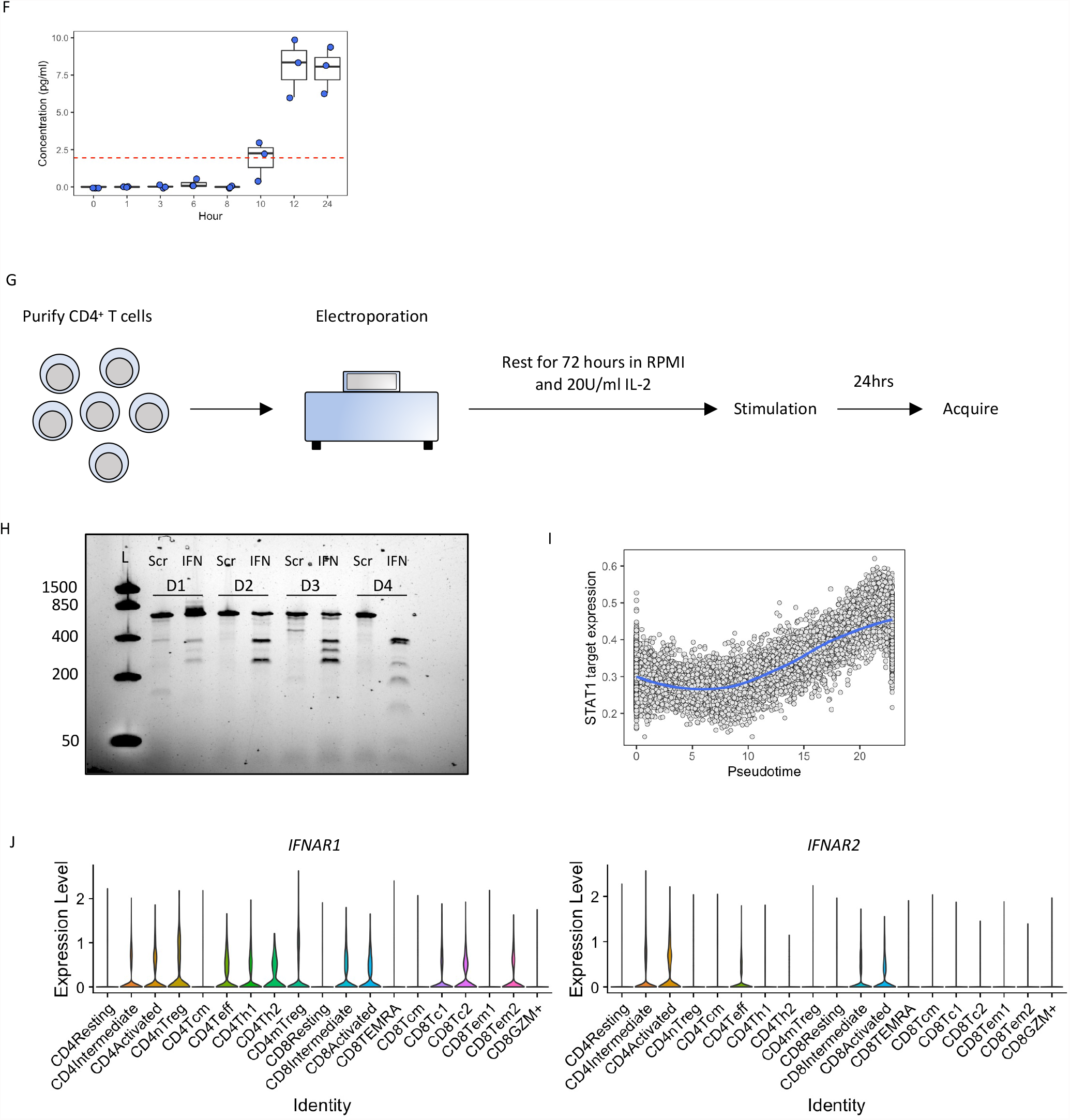
CRISPR mediated knockdown of IFNA in CD4+ T cells (A) Mean pathway expression change using genes from the gene set ‘Reactome interferon signaling’ in unstimulated versus 24 hour stimulated CD4^+^ T cell subtypes. (B) IFNG expression across naïve CD4^+^ T cell populations (C) IFNγ expression in naïve (CD45RA^+^) and memory (CD45RO^+^) T cells (D) Qvals of type 1 interferon signatures (red dots) in T cells after activation (E) Boxplots show expression of all IFNA genes detected in each public dataset. Data were taken from each indicated repository, normalized, and filtered to show all IFNA genes present. (F) IFNα production by CD4^+^ T cells upon activation. CD4^+^ T cells were stimulated using anti CD3 + anti CD28 for the indicated time, and cell supernatants were measured for the presence of IFNα (G) Outline of CRISPR protocol. Cells were stimulated using 2ug/ml anti-CD3 + anti-CD28 (H) T7 assay for analysis of CRISPR efficiency. WT band of IFNA is expected at 589, with fragments in the CRISPR-Cas9 edited samples expected to be at: 346, 302, 284, and 240. L = ladder, Scr = scrambled, IFN = IFNA1 CRISPR-Cas9 targeted deletion (I) Average expression of STAT1 targets over naïve T cell activation, calculated through mean gene expression per cell. STAT1 targets were taken from the STAT1_01, STAT1_02, and STAT1_03 gene sets available on MSigDB. (J) Expression of IFNAR1 and IFNAR2 across T cell subsets. Subsets correspond to those highlighted in the UMAP in Figure 2B.

**Supplementary Figure 6.**
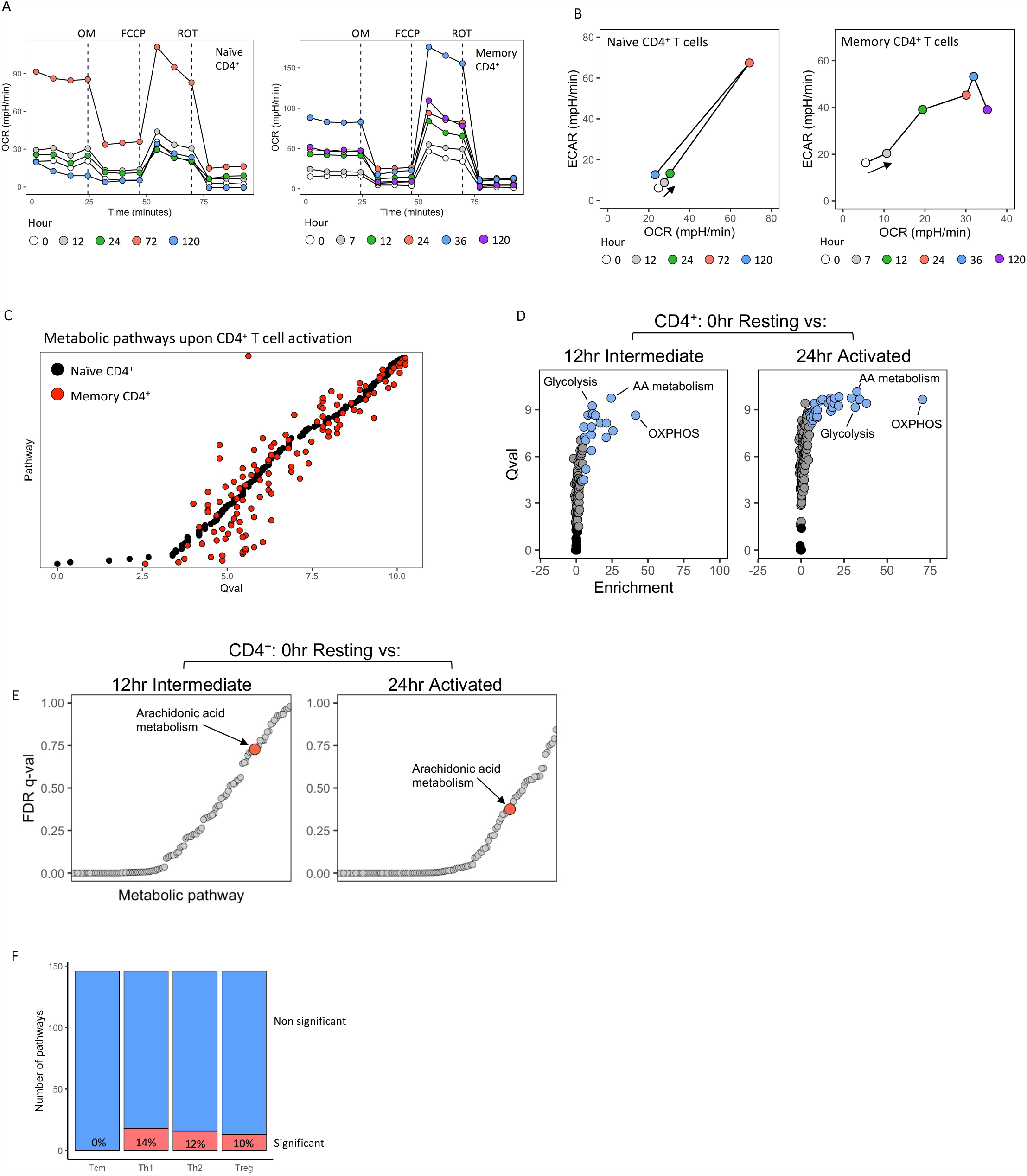
Metabolic analysis of CD4+ T cell activation (A) Seahorse analysis of naïve (left panel) and memory (right panel) CD4^+^ T cells. Cells were activated with anti-CD3 and anti-CD28 for the indicated amount of time, after which, oxygen consumption rate (OCR) was measured. OM = oligomycin, FCPP = carbonyl cyanide-p-trifluoromethoxyphenyl-hydrazon, ROT = rotenone. (B) Seahorse analysis of naïve (left panel) and memory (right panel) CD4^+^ T cells.. Cells were activated with anti-CD3 and anti-CD28 for the indicated amount of time, after which, oxygen consumption rate (OCR) and extracellular acidification rate (ECAR) was measured. (C) Qvals for metabolic pathways upon naïve and memory CD4^+^ T cell activation. SCPA was used to assess pathway distribution changes in naïve and memory CD4+ T cells over 0 and 24 hours of stimulation. (D) Volcano plot showing Qval output from SCPA plotted against pathway enrichment at 12 and 24 hours post stimulation. Black points show non significant pathways, gray points show significant pathways with no enrichment, and blue points show significant pathways that also show enrichment. (E) Gene set enrichment analysis of metabolic pathways in naïve CD4+ T cells. Metabolic pathways in 0hr resting CD4^+^ T cells were compared to 12hr (left panel) and 24hr (right panel) stimulated CD4+ T cells, with arachidonic acid metabolism highlighted in red. (F) Number of significantly perturbed metabolic pathways in resting naïve cells versus resting memory population indicated. Significance was classed as a Qval > 5. Percentages indicate the proportion of significant pathways

**Supplementary Figure 7.**
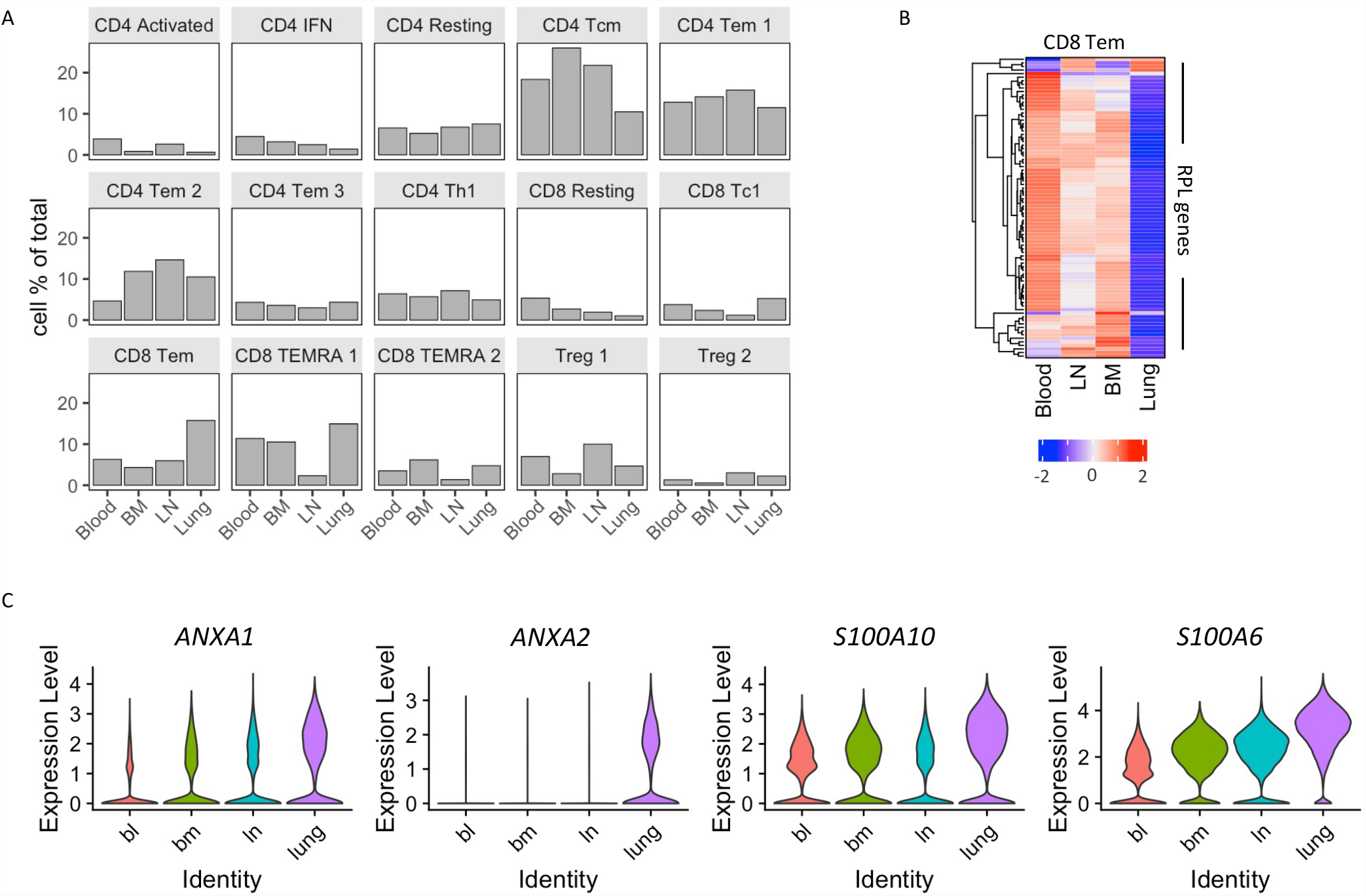
Characterizing T cells across tissue sites (A) Proportional frequencies of T cell lineages across tissue sites (B) Heatmap of ribosomal gene family member expression across tissue sites in CD8^+^ Tem cells (C) Expression of prostaglandin family genes across tissue sites

